# A naturally evolved mutation in cavefish *mc3r* reshapes energy homeostasis across development in a nutrient-limited environment

**DOI:** 10.64898/2026.09.28.753823

**Authors:** Tyler Guzman, Tathagata Biswas, Vikas Tiwari, Edward Thum, Kyle Weaver, Kym Delventhal, Jonathan Wren, Cassidy Petree, Gaurav K. Varshney, Michael C. Rudolph, Chris Seidel, Nicolas Rohner, Jaya Krishnan

**Author notes:** Department of Anthropology, Baylor University, Houston, Texas, USA.

## Abstract

Food availability in natural habitats is often unpredictable and drives the selection of adaptive strategies in organisms to survive food scarcity. However, the genetic and molecular mechanisms that enable animals to survive under such nutritional extremes are poorly understood. Naturally adapted organisms therefore provide powerful genetic systems for uncovering the mechanisms that maintain energy homeostasis under nutritional extremes. Here, we leveraged the cavefish *Astyanax mexicanus* to identify a naturally occurring *Thr166Met* missense mutation in the melanocortin 3 receptor (*mc3r*) that reprograms energy homeostasis to facilitate adaptation to nutrient deprivation. To determine its functional consequences, we introduced the mutation into zebrafish *mc3r* using CRISPR/Cas9 genome editing. We show that adult *mc3r* mutants exhibit increased body size, adiposity, and hyperglycemia relative to wild-type animals, recapitulating key metabolic features of Tinaja cavefish and identifying *mc3r* as the first causal gene behind cavefish hyperglycemia. Investigations for early developmental consequences of the mutation revealed accelerated yolk utilization in mutant zebrafish larvae and strikingly elevated metabolic rates relative to WT, thus, uncovering a previously unrecognized role for *mc3r* in regulating energy expenditure. Importantly, the phenotype was also observed in Tinaja cavefish revealing a novel cavefish phenotype and establishing causal role for *mc3r* in elevated metabolic rate phenotype of cavefish. Together, these findings reveal that the *Thr166Met* variant produces distinct physiological consequences across life stages, enhanced energy storage and hyperglycemia in adulthood despite elevated larval energy expenditure. We propose that increased larval metabolism may support the energetic demands of enhanced foraging, whereas increased adiposity and blood glucose later in life provide energy reserves during periods of nutrient scarcity. More broadly, our study demonstrates how a single naturally evolved coding variant can reshape energy homeostasis across development and contribute to adaptation to nutrient-poor environments.

## Introduction

Energy homeostasis is the ability of an organism to dynamically coordinate energy acquisition, allocation, storage, and expenditure in response to changing energetic demands and nutrient availability ^1,2^. Maintaining this balance is essential for growth, reproduction, survival, and adaptation across diverse ecological environments ^3^. Species that have evolved under extreme nutrient conditions provide powerful opportunities to identify naturally selected genetic variants that regulate these processes ^4^. The Mexican cavefish, *Astyanax mexicanus*, exists as river-dwelling surface fish and several independently evolved cave populations inhabiting the caves of northeastern Mexico ^5^. These perpetually dark, nutrient-limited caves lack primary producers and conventional food webs, exposing cavefish to prolonged periods of food scarcity punctuated by episodes of abundance following seasonal floods ^5^. Thus, unlike the surface fish, cavefish have evolved to thrive under fluctuating nutritional availability.

Adaptation to fluctuating nutrient availability requires mechanisms that translate nutritional and hormonal signals into coordinated changes in feeding, energy expenditure, nutrient utilization, and energy storage. G protein-coupled receptors (GPCRs) are important for mediating such changes because they couple extracellular signals to intracellular pathways that regulate whole-organism physiology ^6–9^. Consistent with this idea, coding variation in two members of the melanocortin receptor family of GPCRs, *mc1r* and *mc4r*, have already been implicated in cavefish adaptation, suggesting that GPCRs may represent a recurrent target of evolutionary change ^10,11^. We therefore performed a GPCR-wide screen for coding variants that could contribute to metabolic adaptation in cavefish. This analysis identified a non-conservative *Thr166Met* amino acid substitution in a highly conserved region of melanocortin 3 receptor (*mc3r*) in the Tinaja cavefish. Mc3r is a regulator of nutrient partitioning and energy expenditure, but the distinct function of this gene has been less clear as *mc3r* null rodents have provided phenotypes that often overlapped with those of *mc4r* null rodents ^6,12–15^. However, the fixation of missense variants in both *mc4r* and *mc3r* in the Tinaja cavefish indicate distinct roles for the two genes in metabolic adaptation ^10^. Hence, we took that as an opportunity to functionally characterize the role of *mc3r* and understand its relevance in metabolic adaptation in cavefish.

To determine the functional consequences of the cavefish *mc3r* variant, we generated a zebrafish CRISPR knock-in carrying the *Thr166Met* substitution. Taking advantage of the external development of both zebrafish and *Astyanax*, we uncovered a previously unrecognized role for *mc3r* in developmental metabolism. In adults, the *Thr166Met* mutation was sufficient to recapitulate key metabolic features of Tinaja cavefish, including increased body size, adiposity, and hyperglycemia, identifying *mc3r* as the first gene causally associated with the cavefish hyperglycemic phenotype. We also observed increased metabolic rates in the mutation zebrafish larvae. Importantly, the elevated metabolic rate was also observed in Tinaja cavefish revealing a novel cavefish phenotype and establishing a causal role for *mc3r* in cavefish. Together, we propose that increased larval metabolism may support the energetic demands of enhanced foraging, whereas increased adiposity and elevated blood glucose later in life provide energy reserves during periods of nutrient scarcity. Together, these findings reveal *mc3r* as a regulator of life-stage-specific energy homeostasis and suggest that altered melanocortin signaling contributes to the metabolic strategies that enable cavefish to persist under fluctuating nutrient availability. More broadly, our study demonstrates how a single naturally evolved coding variant can dynamically reshape metabolism across development and contribute to adaptation in nutrient-poor environments.

## Results

### *mc3r* is mutated in cavefish

G protein-coupled receptors (GPCRs) are central regulators of energy homeostasis, translating environmental cues into coordinated metabolic responses ^7–9,16^. We therefore reasoned that naturally selected variation in energy homeostasis-associated-associated GPCRs could contribute to the remarkable metabolic adaptations of cavefish. To identify naturally selected mutations in GPCRs underlying metabolic adaptation, we performed an in-silico screen of metabolism-related GPCRs using a published multi-genome dataset ^17^ (See **Supplementary Data S1** for detailed methods). Using a custom SnpEff database, we identified coding variants in 54 GPCRs **(Supplementary Data S1)**. Among these, we observed a novel *Thr166Met* missense mutation in the melanocortin 3 receptor (*mc3r*) in the Tinaja cavefish population **(Figure 1A)**. Notably, other members of the melanocortin receptor family of proteins have repeatedly emerged as targets of adaptive evolution: *mc1r* underlies pigmentation loss in cavefish and beach mice, whereas *mc4r* mutations promote hyperphagia in domestic pigs and several cavefish populations ^10,11,18–20^. Although studies in mammals have demonstrated that *mc3r* and *mc4r* regulate overlapping aspects of energy homeostasis including body mass regulation and nutrient allocation into muscle and fat, the physiological functions of *mc3r* remain substantially less well defined than those of *mc4r* ^6^. The independent selection of missense variants in both *mc3r* and *mc4r* within the Tinaja population suggested that these seemingly closely related receptors could make distinct contributions to its metabolic adaptation. We confirmed the presence of the *Thr166Met* mutation by sequencing the *mc3r* (ENSAMXG00000026088; Chr6:1997695-2001170) gene from multiple cavefish populations and noted that the Threonine to Methionine missense mutation (*Thr166Met*, corresponding to *C497T* nucleotide variation) was unique to Tinaja cavefish Mc3r. The other cavefish populations including Pachòn and Molino had the reference surface fish allele **(Supplementary Data S2)**. *Thr166* is highly conserved across vertebrates, including human, mouse, zebrafish, surface fish, and Pachón cavefish, whereas the Tinaja lineage uniquely carries a methionine substitution at this position (Figure 1A). The other variants discovered within the *mc3r* gene were present in less conserved regions of the protein **(Supplementary Data S2)**. Thus, together with its evolutionary conservation and population specificity, these observations prioritized *Thr166Met* for functional analysis.

**Figure 1:**
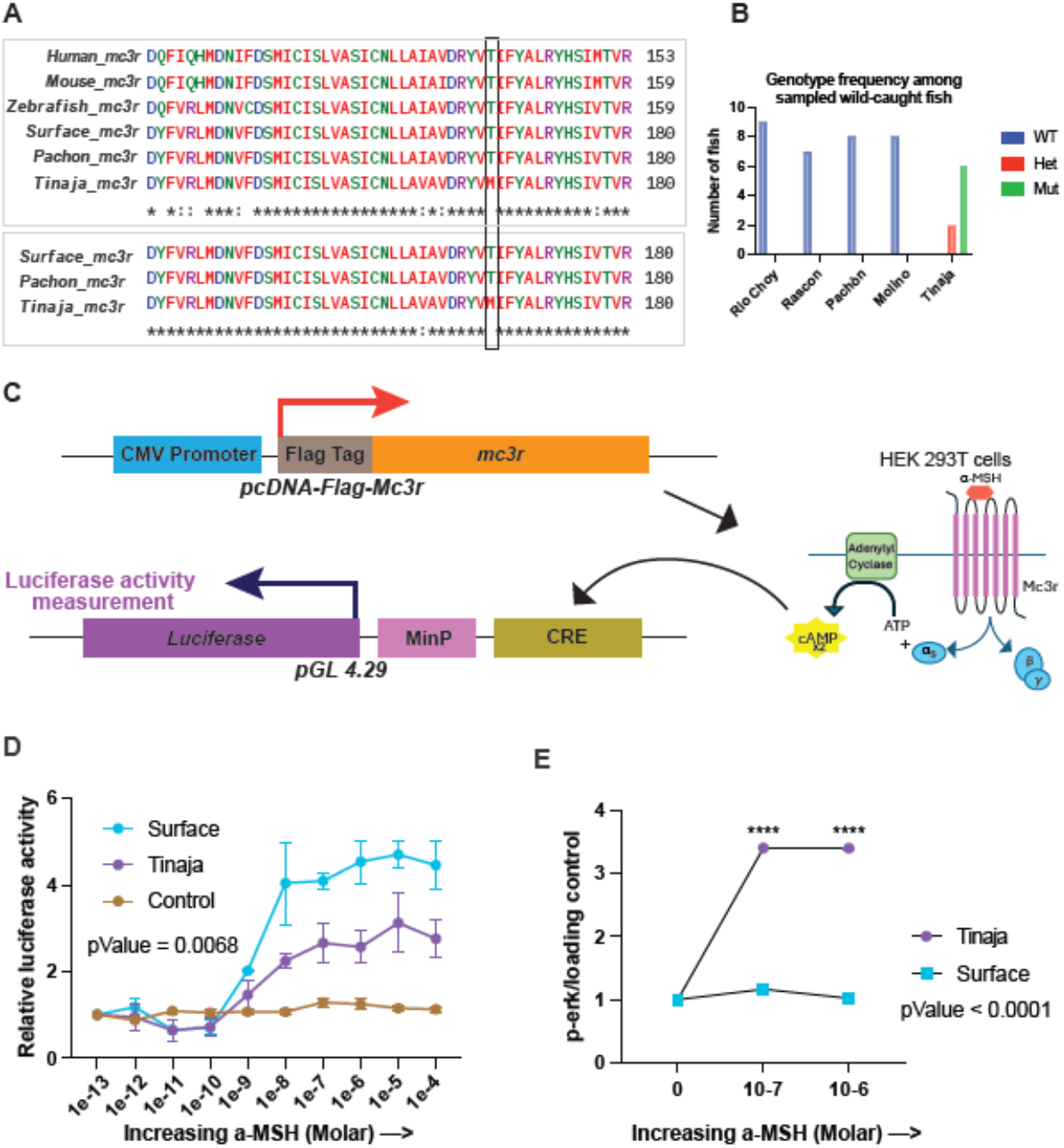
A novel *Thr166Met* mutation in Tinaja Melanocortin 3 Receptor: **(A)** Sequence alignment of partial mc3r proteins showing the *Thr166Met* mutation (Black box). The upper panel shows alignment across multiple species. The lower panel show alignment between morphotypes of *Astyanax mexicanus***. (B)** Genotyping results for the *Thr166Met* mutation from multiple wild-caught *Astyanax* populations. **(C)** Schematic for the ligand binding assay performed to test the effect of the mutation on the cAMP signaling pathway. **(D)** Results from the ligand-binding assay showing Tinaja Mc3r has lower cAMP signaling activity than Surface fish Mc3r when stimulated with the ligand alpha-melanocortin stimulating hormone (a-MSH). **(E)** Relative densitometry quantification of western blot showing p-Erk signal is higher in HEK293T cells transfected with Tinaja-Mc3r in comparison to cells transfected with Surface-Mc3r upon stimulating with the ligand a-MSH.

A *Thr166Met* mutation causes a non-conservative change of a polar amino acid to a sulphur containing non-polar amino acid that can alter protein stability. The *Thr166Met* mutation in cavefish Mc3r lies in a highly conserved region of the protein **(Figure 1A)** responsible for receptor activation. Specifically, a mutation in the adjacent Isoleucine is implicated in human obesity ^21^. Importantly, the mutation was present in all the 8 Tinaja fish that were sequenced in the earlier study (meta-analysis of ^17^) with 6 out of 8 Tinaja fish to be homozygous and 2 heterozygous for the mutation **(Figure 1B)**. None of the other cavefish populations had the mutation in heterozygous or homozygous condition. This hinted toward the positive selection of the mutation and a probable selective advantage the mutation could offer the Tinaja cavefish in the wild. Lastly, several mutations in this region of the *mc3r* have been implicated in altered signaling and metabolic disease in human patients as well indicating its important role in modulation of organismal physiology and likely that the *Thr166Met* mutation could alter cavefish physiology ^13^.

### The *Thr166Met* missense mutation reduces the activity of melanocortin 3 receptor

Melanocortin signaling functions via cAMP signaling and Erk signaling pathways ^22^. To directly test the effect of the *Thr166Met* mutation on its cAMP signaling activity, we performed a ligand binding assay **(Figure 1C)**. We compared the response of surface fish Mc3r and Tinaja cavefish Mc3r to one of its ligands alpha melanocortin stimulating hormone (α-MSH) in a heterologous system ^12^. The coding sequences of surface and Tinaja *mc3r* were tagged with FLAG epitope and cloned in overexpression vectors and co-transfected into HEK293T cells with a plasmid containing luciferase cloned downstream to cAMP response element (CRE) **(Supplementary Data S3,4).** The melanocortin signaling was induced by stimulating the cells with α-MSH that in turn produced cAMP. cAMP-induced luciferase expression served as a measure of signaling through Mc3r. We observed that luciferase expression levels increased in both, cells expressing surface fish *mc3r* and the Tinaja *mc3r*. However, the maximal activity of Tinaja Mc3r obtained with high concentrations of ligand MSH were significantly lower than that of surface fish Mc3r (**Figure 1D**). Next, to compare the surface and Tinaja Mc3r protein function on the downstream ERK signaling, we stimulated the HEK cells overexpressing surface fish or Tinaja *mc3r* with increasing concentrations of the α-MSH ligand and checked for phospho-Erk levels using western blotting **(Figure 1E, Supplementary Figure S1)**. We observed that the *Thr166Met* mutation enhances activity of the downstream ERK signaling pathway . This was in line with the predicted opposing effects these pathways downstream of *mc3r* have ^22^. Thus, we predicted that the *Thr166Met* mutation can potentially lead to diverse phenotypic alterations. Thus, we hypothesized that the reduced melanocortin signaling through Tinaja Mc3r could significantly affect Tinaja cavefish physiology.

### Development of a *mc3r^Thr^*^166^*^Met^* mutant zebrafish line

Decreased melanocortin signaling can have broad effects on the physiology of an organism ^6,23^. However, as there are numerous other genetic variations between surface fish and Tinaja cavefish, the phenotypic dissection of the *Thr166Met* mutation in *Astyanax* is highly constrained. Thus, we leveraged the zebrafish model system and generated a precise *Thr166Met* mutation in zebrafish using CRISPR/Cas9-based homologous recombination method. To obtain precise genome editing, we also introduced silent mutations nearby PAM sites in the donor DNA that prevented the modified genome from being targeted by the gRNA again **(Supplementary Figure S2, Supplementary Data S5)** ^24^. Homozygous mutant lines were created and compared with wild type (WT) zebrafish for the purpose of this study. This complementary zebrafish model provided the power to inform how the specific *Thr166Met* mutation shapes the physiology of cavefish.

### *mc3r^Thr^*^166^*^Met^* mutant adults are obese and hyperglycemic

We began the phenotypic analysis of the homozygous zebrafish mutants by measuring the standard lengths of the adults. We observed that only mutant males were marginally longer **(Figure 2A)**, while the mutant and WT females did not show any significant differences in their standard lengths **(Figure 2B)**. Next, we observed that the mutants were heavier than the WT zebrafish. Both male and female mutant zebrafish adults weighed more than the WT indicating no sexual dimorphism in the trait **(Figure 2C, D)**. The higher weights of the mutant adults prompted us to test if the increased weight signified obesity phenotype. EchoMRI analysis revealed a striking increase in fat mass in mutant fish compared with WT controls **(Figure 2E–H)**, a phenotype observed in both males and females. Mutant fish also exhibited higher absolute lean mass than WT fish. However, when normalized to body weight, lean mass constituted a smaller proportion of total body mass in mutants, indicating that the increase in fat mass outpaced the increase in lean mass. Together, these data demonstrate that the mutants exhibit elevated adiposity characterized by increased fat accumulation and a reduced relative contribution of lean tissue **(Figure 2I–L)**.

**Figure 2:**
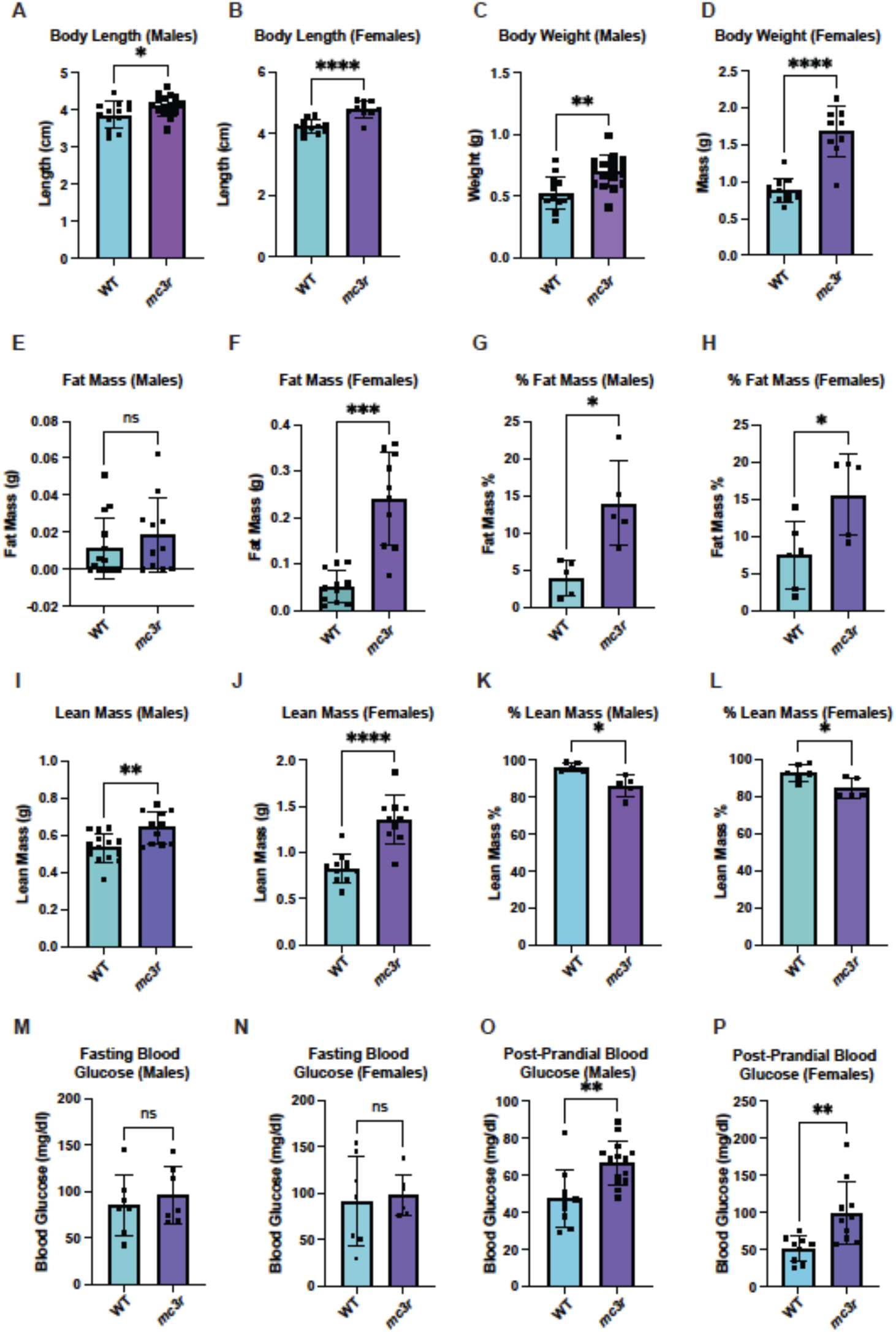
Phenotypic characterization of *Thr166Met* zebrafish mutant adults: Wild-type (WT) and mutant (*mc3r*) adult males and females were compared for various phenotypes including: Standard body length **(A, B)**; Body weight **(C, D)**; Fat mass **(E, F)**; % Fat mass **(G, H)**; Lean mass **(I, J)**; % Lean mass **(K, L)**; Fasting blood glucose **(M, N)**; Post-prandial blood glucose **(O, P)**. n for males = 13, 17. n for females = 11, 10. P values were calculated using Two-tailed Welch’s test * < 0.05, ** < 0.01, **** < 0.0001, ns = not significant.

As obesity often is accompanied by hyperglycemia ^25^, we tested for blood glucose levels in the WT and *mc3r^Thr^*^166^*^Met^* mutant adults. We observed little or no difference in the typical fasting glucose levels between the genotypes **(Figure 2M, N)**. In contrast, post-prandial blood glucose levels measured one hour after feeding were significantly elevated in *mc3r^Thr^*^166^*^Met^*mutants relative to WT controls, indicating impaired glucose tolerance following nutrient intake. **(Figure 2O, P)**. These findings are notable in the context of previous studies of cavefish metabolism. A previous study ^26^ reported that elevated fasting blood glucose in surface × Tinaja F2 hybrids is a complex, non-Mendelian trait but found no significant association between blood glucose levels and the cavefish *insra^P^*^211^*^L^* mutation, although all individuals exhibiting the most extreme glucose values carried the mutation. This suggested that the *insra* variant may contribute to, but is not sufficient for, altered glucose regulation. In contrast, introduction of the cavefish *mc3r^Thr^*^166^*^Met^* allele into zebrafish was sufficient to produce a robust post-prandial hyperglycemia phenotype, providing direct functional evidence that a naturally occurring cavefish mutation can alter glucose tolerance. To our knowledge, this represents the first cave-derived variant for which a causal role in dysregulated glucose metabolism has been experimentally demonstrated in vivo.

### Transcriptomic analysis identifies pathways altered in *mc3r^Thr^*^166^*^Met^* mutants

Next, to elucidate how altered Mc3r signaling affects development, we took an unbiased transcriptomic approach. WT and mutant 14dpf larvae were collected and bulk RNA-seq was performed in 3 biological replicates. We chose 14dpf specifically to get the transcriptomic signature at the whole organism level and as most organs are developed by this stage including the adipose tissue ^27^. We obtained an average of 45 million reads for each sample and performed differential gene expression analysis. The PCA plot revealed that the transcriptomic profiles of *mc3r^Thr^*^166^*^Met^*were significantly affected as the x-axis has a clear separation of genotypes **(Figure 3A)**. The y-axis reflects sample-to-sample variation, and no unexpected sub-clustering was observed, indicating overall consistency among biological replicates. For differential gene expression (DGE) analysis, stringent thresholds were applied (padj < 0.01 and |log2FC| > 0.585) because a large number of genes were significant at lower cutoffs; these stricter criteria reduced false positives while retaining biologically meaningful expression changes **(Supplementary Data S6)**. There were 1456 genes that were significantly upregulated (padj < 0.01, log2FC > 0.585) and 781 genes were significantly downregulated (padj < 0.01, log2FC < -0.585) in *mc3r^Thr^*^166^*^Met^* compared to the WT **(Supplementary Data S7, 8) (Figure 3B)**.

**Figure 3:**
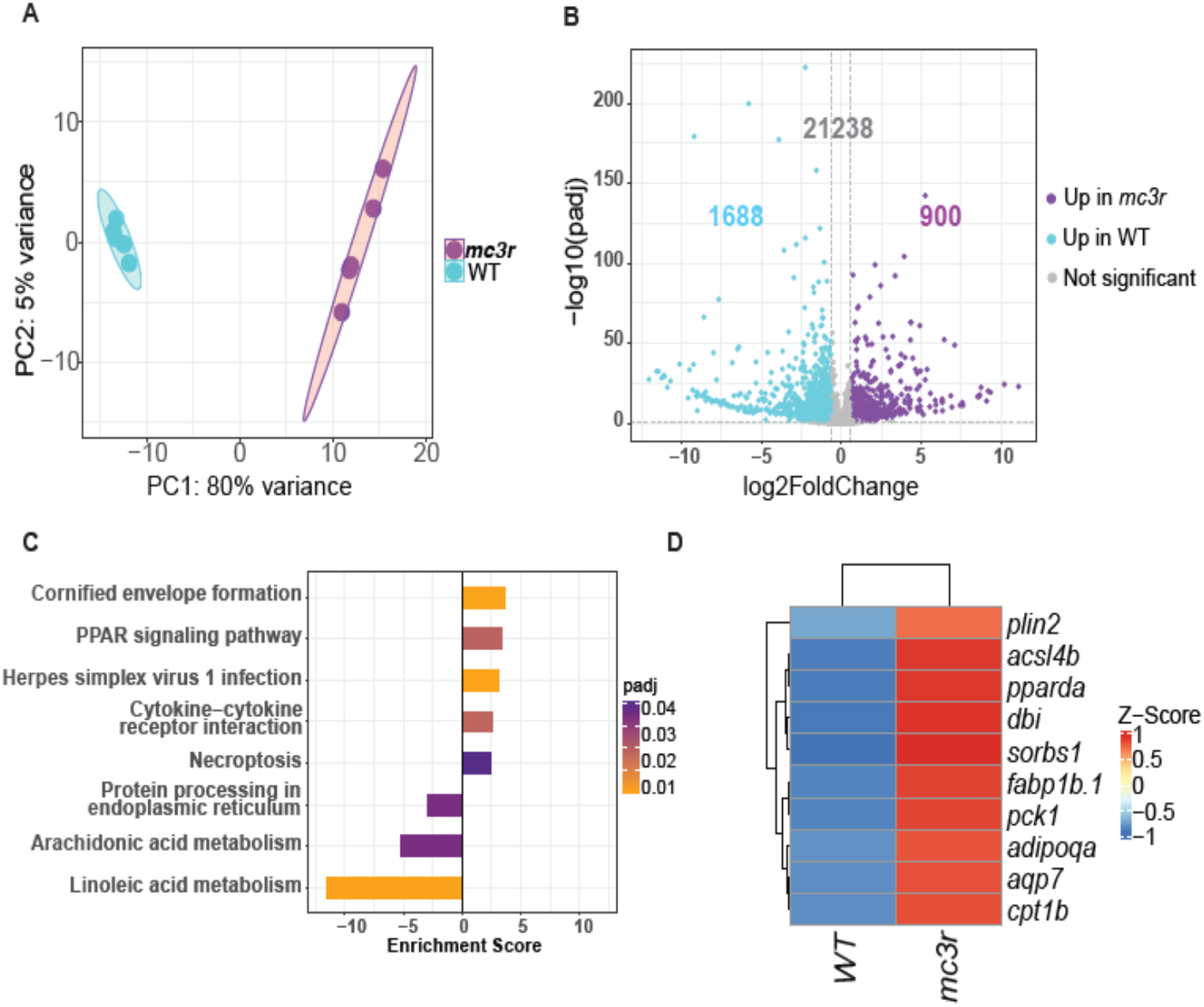
Transcriptomic analysis of 14dpf wild-type and mutant larvae. **(A)** PCA plot showing the comparative distribution of samples in the dataset. **(B)** Volcano plot showing the differentially expressed genes. **(C)** KEGG pathway analysis showing pathways up-regulated (enrichment score > 0) and down-regulated (enrichment score < 0)in the mutants. Enrichment score is the ratio of differentially expressed genes within a pathway compared to the expected background. **(D)** Heatmap showing expression levels of genes under the PPAR signaling pathway.

Using the differentially regulated genes, a KEGG pathway analysis was done to determine the dysregulated pathways **(Figure 3C)**. The two main categories that were enriched were the immune-related pathways as well as the metabolic-related pathways. Among the immune system related pathways, we obtained ‘Herpes simplex virus infection’ and ‘Cytokine receptor interaction’. For the metabolic pathways, we obtained ‘Steroid biosynthesis’, ‘Metabolism of xenobiotics’, ‘Linoleic acid metabolism’, ‘Arachidonic acid metabolism’, and ‘PPAR signaling pathway’ **(Figure 3C, Supplementary Data S9)**.

### *mc3r^Thr^*^166^*^Met^* mutants have a higher metabolic rate

A closer look at the PPAR signaling pathway enriched among the differentially expressed genes, we observed the upregulation of *pparδa, fabp1b.1* and *cpt1b* **(Figure 3D)**. Increase in *pparδ, fabp1b.1* and *cpt1b* causes increase in fatty acid oxidation ^28–30^. This came across as an unexpected and a paradoxical observation. We originally predicted that the transcriptome would support enhanced fat synthesis rather than breakdown to cause the obesity phenotype in the adults. Thus, an increase in fatty acid oxidation would translate to an altered (or increased) metabolic rate. To test this, we performed an Alamar blue assay. Alamar Blue assay estimates metabolic rate by measuring the cellular redox metabolism via the reduction of resazurin to the fluorescent resorufin ^31^. We observed remarkably higher metabolic rates in the *mc3r^Thr^*^166^*^Met^* mutants compared to WT at larval stages **(Figure 4A)**. We also measured if the high metabolic rate continues even in adult stages by repeating the Alamar Blue assay with mutant and WT adults **(Figure 4B)**. Interestingly, the mutants continued showing higher metabolic rates than the WT even as adults. The difference was retained even when difference in their body mass was accounted for **(Supplementary Figure S3A)**, thus establishing a novel role for *mc3r* in vertebrate metabolism.

**Figure 4:**
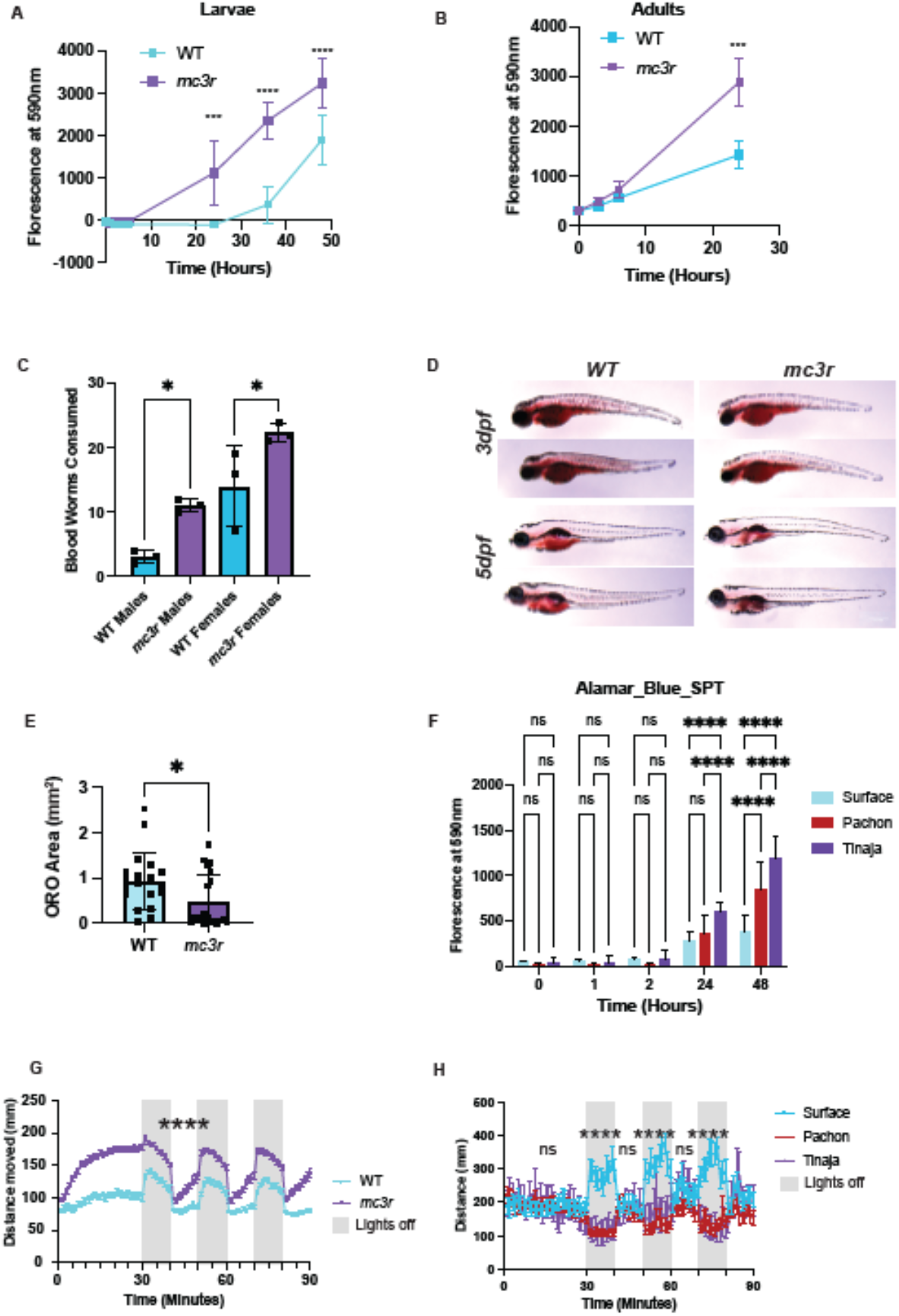
*mc3r^Thr^*^166^*^Met^* mutants have higher metabolic rates than WT. **(A)** Alamar Blue assay showing metabolic rates are higher in the mutants than WT. Assay was done on freshly hatched 3dpf larvae n = 13, 13. P value < 0.0001 using 2way ANOVA. **(B)** Alamar Blue assay showing metabolic rates are higher in the mutant adults relative to WT. Assay was done on 15mpf fish. n = 10, 10. P value < 0.001 using a custom linear mixed model. **(C)** Feeding assay done on WT and mutant zebrafish juveniles. n = 15, 13. P Value < 0.05 using T-test. **(D)** Brightfield images of 3dpf and 5dpf larvae stained with Oil Red O showing that yolk consumption is higher in the mutants than the WT **(E)** Quantification of Oil Red O staining in figure 4B. n = 19, 20. P value <0.05 using 2-tailedT-test **(F)** Alamar Blue assay of Surface, Pachòn, and Tinaja larvae. The assay was done on 3dpf larvae. n = 24, 24, 17. P values calculated using 2way ANOVA. **(G)** Locomotion/behavior assay done on 5dpf mutant and WT zebrafish larvae. n = 48, 48. P < 0.0001 using Kruskal-Wallis test. **(H)** Locomotion/behavior assay done on 8dpf. Surface, Pachòn, and Tinaja larvae. n = 16, 15, 10. P values using Kruskal-Wallis test. P value * < 0.05, ** <0.01, *** <0.001, ****<0.0001.

### *mc3r^Thr^*^166^*^Met^* mutants have a higher food intake

The next question was how the *mc3r^Thr^*^166^*^Met^* mutants with higher metabolic rates become obese and hyperglycemic adults. One plausible explanation is that they compensate for increased energy demand by increasing food intake. We measured the amount of food the fish ate and observed a substantial increase in the number of worms they consumed, potentially explaining the transition of the mutants from fast yolk consumers to high adipose storers **(Figure 4C)**. We measured standard length and width (Height of the Body at the Anterior Margin of the Anal Fin or HAA) at 7dpf, 14dpf, and 21dpf and found no significant differences between the mutant and the WT suggesting that altered growth rate during early development is not a factor influencing the increased metabolic rate or higher appetite in the mutants **(Supplementary Figure S3B)**. We also observed the consumption of the maternal yolk deposits in larvae. Using oil red O staining, we visualized the yolk sizes and observed that while, right after hatching, at 3dpf the yolk sizes looked similar, by 5dpf the mutants had exhausted most of it again indicating higher nutrient consumption **(Figure 4D, E)**. We ruled out any developmental stage differences between the mutants and the WT by ensuring similar embryonic stages in the beginning of the experiment **(Supplementary Figure S3C).**

### Cavefish larvae have a higher metabolic rate

We next asked if the phenotype observed in the zebrafish mutant are also seen in Tinaja. Earlier studies have already reported higher fat-to-lean mass ratio, hyperphagia, and glucose intolerance in Tinaja as compared to surface fish adults ^10,26,32,33^. We tested if the novel phenotype of higher metabolic rate that we observed in the zebrafish *mc3r* mutants would also be seen in the Tinaja population. Using the Alamar Blue assay, we quantified cellular reducing capacity as a proxy for metabolic rate and indeed, as compared to surface fish, the Tinaja larvae had remarkably higher metabolic rate **(Figure 4F)**. This clearly demonstrated the causal role of *mc3r* mutation in the increased metabolic rate of cavefish. Interestingly, even Pachòn larvae showed higher metabolic rate than surface fish larvae at the 48h timepoint, although it was still significantly lower than that of Tinaja larvae **(Figure 4F)**. This hints at the possibility that metabolic rate is a complex genetic trait and that additional mutations/genes beyond *mc3r* regulate the trait. It needs to be noted that although the cavefish larvae show elevated metabolic rates, previous studies with Pachòn cavefish have shown that cavefish adults have lower metabolic rate than adult surface fish. This may mean that metabolic rate regulation is morphotype and/or developmental-stage-specific and is a topic for future investigation.

### Higher metabolic activity supports increased physical activity *mc3r^Thr^*^166^*^Met^* mutants

We sought to understand why elevated metabolic activity during larval stages would be favored in an environment characterized by limited and fluctuating nutrient availability. The *mc3r^Thr^*^166^*^Met^* mutation is found in the Tinaja cavefish population. The Tinaja cave has a unique geology and allows rainwaters to flood the caves every year during the monsoons. After the monsoons, the water recedes creating a seasonal cycle of high and low water levels. The fluctuating water levels would also fluctuate the density of the food present in the water requiring increased foraging activity to find food before the yolk deposits finish. Thus, we reasoned that the elevated metabolic rates could be to support increased locomotor or physical activity. To test this, we turned back to our zebrafish lines. We quantified physical activity using the Noldus Zebrafish behavior monitoring system and observed that the mutants on an average had more activity than the WT **(Figure 4G)**. As physical activity can be influenced by vision, we measured movement in alternate periods of 10 min of light and dark and obtained a similar enhanced activity of mutants in both light and dark conditions. **(Figure 4G)**. We confirmed our results by video recording individual larvae and tracing their movement using Zebra Zoom tool **(Supplementary Fig. S4)** thus together explaining the faster yolk consumption the mutants relative to WT.

We also measured locomotion in surface fish and Tinaja and Pachòn cavefish larvae and observed no significant difference between surface and cave morphotypes in light ON conditions **(Figure 4H)**. However, in dark, the surface fish increased their physical activity similar what was seen in zebrafish mutant and WT **(Figure 4H)**. Such increase in activity in dark has been reported as a standard response to light in zebrafish and by extension in surface fish that is likely not a consequence of the mutation ^34^. The absence of elevated locomotor activity in Tinaja larvae relative to surface fish suggests that the behavioral effect of the *mc3r Thr166Met* variant is modified by the broader cavefish genetic background. Such genetic background dependence could reflect epistatic interactions with other cave-adapted alleles affecting locomotor behavior. Thus, although locomotor activity is not elevated in Tinaja larvae, the zebrafish mutant provides causal evidence that altered *mc3r* signaling is sufficient to increase activity.

## Discussion

Survival in nutritionally unpredictable environments depends on the ability to dynamically balance energy expenditure with acquisition and storage, making energy homeostasis a critical target of adaptation. This challenge is particularly relevant in *Astyanax mexicanus* cave populations, which experience marked differences in the timing and availability of food resources. In some caves, such as Pachón, bat populations provide an indirect source of nutrients through their excrements. In contrast, caves such as Tinaja experience seasonal flooding that alter both water levels and food availability, exposing Tinaja cavefish to recurrent transitions between nutrient abundance and scarcity. Such conditions are likely to impose strong selective pressure for flexible energy homeostasis mechanisms that allow animals to rapidly adjust energy acquisition, utilization, and storage in response to changing nutritional conditions.

As GPCRs are central to energy homeostasis and response to nutrients ^35,36^, in this study, we screened for coding mutations in metabolism-related GPCRs in the cavefish genome. The screen identified a novel mutation in melanocortin 3 receptor (*mc3r*) that alters energy homeostasis to enable *Astyanax mexicanus* to survive in nutrient-poor cave habitats. Melanocortin signaling is central to metabolic regulation and maintaining energy homeostasis in vertebrates ^23,37^. Specifically, melanocortin 3 receptor regulates the proportion of nutrients that are stored as fat in the body to maintain energy reserves ^12^. The missense mutation (*Thr166Met*) identified in this study occurs within an evolutionarily conserved region of *mc3r* that alters the receptor functionality by decreasing downstream cAMP signaling and increasing Erk signaling ^13^. The mutation was also present in the wild-caught populations of cavefish ruling out the possibility of the mutation being specific to lab-bred populations and supporting its positive selection in the wild.

To test for the phenotypic consequences of the *Thr166Met* mutation, we introduced the mutation into the zebrafish genome and generated a mutant line and performed detailed characterization. We observed the adult males and females to be significantly larger and heavier that the WT confirming that the single amino change in *mc3r* can lead to physiological phenotypes. We also observed that the mutants had significantly higher fat to lean mass ratio than the WT. The high adiposity was also accompanied by higher post-prandial blood glucose, together, recapitulating key metabolic features of Tinaja cavefish and providing a genetic basis for these phenotypes. Notably, *mc3r* is the first reported gene to be causally associated with the hyperglycemia phenotype of cavefish. For the cavefish, this would translate to an adaptive feature for storing ample energy reserves in a nutrient-scarce environment. This was also in line with observations of higher adiposity in *mc3r* mutant mice and humans with *mc3r* mutations ^13,38^.

Investigation of the transcription programs altered in the zebrafish *mc3r^Thr^*^166^*^Met^*larvae revealed the upregulation of *pparδ, fabp1b.1,* and *cpt1b,* genes that are known to enhance catabolism and energy expenditure ^28–30^. Consistent with this observation, our whole-body metabolic rate assay confirmed strikingly higher metabolic rates for the mutants relative to WT. This was also supported by the faster yolk consumption in the mutants as observed using histological staining. Interestingly, despite the high metabolic expenditure, the *mc3r* mutant zebrafish adults develop higher body mass and accumulate higher fat mass by significantly increasing its food intake and compensating for the increased energy demand. Together, these molecular and physiological data identify a previously unrecognized hypermetabolism phenotype associated with the *mc3r Thr166Met* variant and implicate PPAR-linked metabolic pathways as potential mediators of altered energy homeostasis. The recapitulation of elevated metabolic rate in Tinaja larvae further suggests that similar mechanisms may contribute to cavefish metabolic adaptation.

We viewed these observations in the light of cave adaptation and speculated why cavefish larvae would paradoxically have higher metabolic rate in a nutrient limited habitat. Several studies that have in fact studied cavefish have in fact reported a lower energy consumption and metabolic rate. However, all these studies have either focused on adult fish, tissues, or cell lines but not larvae. Our study, therefore, shows for the first time that a mutation in cavefish is associated with higher metabolic rates in the larval stages. This prompted us to ask how such an energetically costly phenotype could confer a selective advantage in Tinaja cavefish larvae. The seasonal increase and decrease in the water levels also alters the density of food available to the fish in the Tinaja cave waters. Adaptation to this situation could happen in two ways: the larvae could forage with less movement to preserve energy and rely on probabilistic availability of food, or they could invest in increased movement and enhanced foraging to increase their chances of finding food. A study on cave salamanders has shown that the ‘sit and wait’ approach may be less beneficial for survival in comparison to ‘move and find’ approach even in nutrient-poor habitats ^39,40^. In line with this hypothesis, we observed increased locomotion in 5dpf mutant zebrafish larvae relative to WT and raised the idea that the enhanced metabolic demand could, in part, be for supporting the elevated physical activity. Upon comparing the locomotory activity levels in Tinaja and surface fish larvae, however, we did not observe any significant difference. It is still noteworthy that the activity levels were similar and not lesser in cavefish relative to surface fish, still supporting the ‘move and find’ rather than ‘sit and wait’ approach for foraging in food-scarce cave habitat. We reason that the lack of elevated locomotion could be due to epistatic masking of the phenotype by other genetic variants in cavefish.

Lastly, the higher metabolic rate phenotype observed in the Pachòn population, which does not harbor the *mc3r* mutation, opens the possibility that the trait could be influenced by other genes as well. Future studies will leverage the ability to make surface x cave and cave x cave hybrids for QTL (Quantitative Trait Loci) analyses to identify other genetic loci controlling the metabolic rate phenotype of various cave morphotypes.

Putting together, our study demonstrates how a single naturally evolved coding variant can reshape energy homeostasis across development and contribute to adaptation to nutrient-poor environments.

## Methods

### *Astyanax* husbandry

*Astyanax mexicanus* are housed in polycarbonate or glass fish tanks on racks (Iwaki Aquatic). The surface fish and the cavefish are maintained under similar laboratory conditions ^41^. They are provided with the same food, and both kept under the same 14/10 h light/dark cycle. Each rack uses an independent recirculating aquaculture system with mechanical, chemical and biologic filtration, and UV disinfection. Water quality parameters are maintained within safe limits (upper limit of total ammonia nitrogen range, 1 mg l^−1^; upper limit of nitrite range, 0.5 mg l^−1^; upper limit of nitrate range, 60 mg l^−1^; temperature set-point, 22 °C; pH, 7.65, specific conductance, 800 μS cm^−1^; dissolved oxygen, >90%). Water changes range from 10% to 15% daily (supplemented with Instant Ocean Sea Salt (Spectrum Brands)). Adult fish are fed two times per day with Zeigler Silver 3mm Slow Sink (Cat # 306540-18-44) and Artemia. Breeders get an additional feed of frozen blood worms (*Hikari*) during the breeding week. IACUC animal husbandry protocol # 24-16 was approved for the Oklahoma Medical research Foundation for the study.

### Zebrafish husbandry

Adult zebrafish (*D. rerio*) of the TU and *mc3r^Thr^*^166^*^Met^* strains were maintained in a recirculating aquaculture system (Aquaneering) at 28.5 ± 0.5 °C on a 14-h light/10-h dark cycle. Water quality parameters were monitored daily and maintained within safe limits: pH 7.2–7.6, conductivity 400–600 µS cm^−1^ and dissolved oxygen >6.0 mg l^−1^. Fish were fed two times daily with a mixture of commercially available dry food (60% EPAC Black (*INVE Aquaculture*), 15% Zebrafeed (*Techniplast*), 25% Otohime B2 (*Reed mariculture*). For breeding, the evening before breeding, male and female fish were separated by a divider in breeding tanks. The dividers were removed the following morning, at the onset of the light cycle, to enable natural spawning. Embryos were collected within 15 min of spawning and maintained in E3 medium (5 mM NaCl, 0.17 mM KCl, 0.33 mM CaCl2, 0.33 mM MgSO₄, 0.0001% methylene blue) at 28.5 °C.

### Cloning

Coding sequence of surface and Tinaja *mc3r* were obtained using PCR amplification from the respective genomic DNA as *mc3r* is a single exon gene. The amplified sequences were cloned into pcDNA3.1 using Gibson assembly. The cloning scheme and final plasmid sequences are provided as supplementary files **(Supplementary Data S3, 4)**.

### Ligand binding assay

Ligand binding assay was performed according to published protocol with some modifications ^42,43^. Briefly, a 500ng mixture of pcDNA-Flag-Mc3r (300ng), cAMP response element-driven luciferase vector pGL4.29 (250ng, Promega), and Renilla luciferase vector pRL-TK (50ng, Promega) was transiently transfected into HEK 293T cells in a 24-well plate for 24 hours. Media was changed to serum-free media for 12 hours followed by induction of melanocortin signaling with various concentrations of NDP-αMSH (M8764; Sigma) and 1-hour incubation. Luciferase activity was measured using the Promega Dual-Glo® Luciferase Assay System (Cat # E2920). The transfection efficiency was normalized using the Renilla Luciferase activity from the pRL-TK vector. The assay was done twice with 3 replicates each time. One out of the two experiments have been presented in Figure 1. Raw Prism file for the other is provided as **Supplementary Data S9**.

### Western Blot

To measure p-Erk levels, immunoblotting was done as per published protocol with minor modifications ^22^. Briefly, HEK-293T cells were seeded at ∼70% confluency in Poly-D-Lysine-coated 24-well plates (A38904-01, Gibco) and allowed to adhere for 12 h in complete medium containing 10% FBS and 1% penicillin-streptomycin. Cells were transfected with pcDNA-Flag-Mc3r (Surface or Tinaja Mc3r) using Lipofectamine 3000 (L3000008, Invitrogen) according to the manufacturer’s protocol. After 6 h, transfection medium was replaced with complete medium. After 24h incubation, cells were serum-starved for 6 h and stimulated with 0, 10^-9^, 10^-8^, 10^-7^, or 10^-6^ M α-MSH (M8764, Sigma) for 5 min. Cells were then washed with ice-cold PBS and lysed in RIPA buffer containing protease and phosphatase inhibitors. Proteins were separated by 10% SDS-PAGE, imaged using stain-free detection, and transferred to PVDF membranes. Membranes were blocked in 5% non-fat milk in TBST and incubated with rabbit anti-pERK1/2 antibody (1:1,000; 4370T, Cell Signaling Technology), followed by HRP-conjugated goat anti-rabbit secondary antibody (1:10,000; G21234, Invitrogen) for 1 h at room temperature. Chemiluminescent signals were detected using Bio-Rad reagents and quantified in Fiji. pERK1/2 levels were normalized to total stain-free protein and expressed relative to the 0M α-MSH control.

### Zebrafish mutant generation

Generation of a *mc3r^Thr^*^166^*^Met^*zebrafish mutant model was performed using CRISPR/Cas9 gene editing technology ^24^. Potential guideRNA target sites were designed using CCTOP ^44^. The target site was selected by evaluating the predicted on-target efficiency score and the off-target potential ^45^ in addition to the proximity of the double-strand break to the desired mutation site. A second guide was designed for dCas9 to bind upstream of the target site at the edge of the homology arm to support HDR repair. To generate the T145M codon change, a single-stranded DNA oligonucleotide (ssODN) donor was designed containing ∼50 nucleotides of homology from the double-strand break site. The ssODN contained mutations for the T145M change and silent codon changes to disrupt guideRNA binding following the repair event. The ssODN was ordered as an Ultramer from Integrated DNA Technologies(IDT). The selected guideRNA was ordered as an Alt-R CRISPR-Cas9 sgRNA from IDT. Ribonucleoprotein (RNP) complexes were formed using sgRNA at 1uM final concentration with IDT Cas9 HiFi v3 protein at 6.6uM final concentration in 15ul. The RNP was formed by incubating at room temperature for 10 minutes. The ssODN was added at 200ng/uL final concentration to the RNP and delivered to Aquatics for injection into zebrafish embryos.

1dpf embryos were screened for the expected mutations by lysing the embryos and amplifying the specific genomic location. A second round of amplification was done to incorporate sample-specific dual barcodes. All amplicons were pooled and size-selected using ProNex Size-Selective Purification System (Promega). Cleaned pools were quantified on a Qubit Fluorometer and then ran on an Agilent Bioanalyzer to check sizing and purity. Purified pools were run on an Illumina MiSeq 2×250 flow cell. The resulting sequence data was demultiplexed and read pairs were joined. On-target indel frequency and expected mutations were analyzed using CRIS.py ^46^. Adult F0 males were screened by lysing sperm and then followed the sequencing procedure described above. After outcrossing potential founders, subsequent generations were screened by fin clips from adult animals.

### RNA sequencing of zebrafish larvae

For RNA-Seq, RNA was isolated from 14dpf larvae using the Qiagen RNeasy kit following the manufacturer’s instruction. Prior to RNA-seq analysis, quality control measures were implemented. Concentration of RNA was ascertained via fluorometric analysis on a Thermo Fisher Qubit 4 fluorometer. Overall quality of RNA was verified using an Agilent Tapestation 4200 instrument. Following initial QC steps, libraries were generated using the Watchmaker RNA Library Prep Kit (Watchmaker Cat#7K0078) according to the manufacturers protocol. Briefly, polyadenylated mRNA molecules were enriched via hybridization to poly-dT probes. The enriched mRNA molecules were then chemically fragmented, and the first strand of cDNA was generated using random primers followed by second strand synthesis and A-tailing. Stubby adapter molecules were ligated followed by indexing PCR using IDT xGen Unique Dual Indexing primers (IDT Cat# 10012162) on an n6Tec iconPCR instrument. During this amplification individual sample wells were monitored via fluorescence and amplification was stopped for each respective well once the PCR reaction exited log phase. Strand selection occurred during this amplification step. Following PCR each library was pooled according to the fluorescence data obtained by the iconPCR. Final library pools were assayed on the Agilent Tapestation 4200 for appropriate size and quantity and were absolutely quantified using qPCR on an ABI QuantStudio6 instrument with NEB Illumina Library Quantification reagents. Sequencing of the libraries was performed on an Illumina NovaSeq X Plus with PE150 reads.

### Mapping of Zebrafish transcriptome and differential gene expression analysis

Raw reads were mapped using Hisat-3n/2.2.1-3n-0.0.3 ^47^ and the genome assembly GRCz11from NCBI was used as a reference. Gene counts were quantified using FeatureCounts from the software package Subread 2.1.1 ^48^. DeSeq2 was utilized to identify differentially expressed genes (DEG) which had a |Log2FC| > 0.585 and a false discovery rate of 0.01 ^49^.

### Gene Ontology and KEGG Analysis

The total differentially expressed gene (DEG) set was divided into two subsets based on log2 fold change to identify genotype-specific biological processes: genes upregulated in mutants (log2FC > 0.585) and genes upregulated in wild type (log2FC < -0.585). Gene annotations based on NCBI IDs were converted to Entrez IDs using BioMart ^50^ to enable Gene Ontology (GO) and Kyoto Encyclopedia of Genes and Genomes (KEGG) enrichment analyses. Functional enrichment was then performed using the enrichGO and enrichKEGG functions in ClusterProfiler ^49^, with a q-value cutoff of 0.05 to define significantly enriched terms and pathways.

### Adult morphometric and blood glucose measurements

Upon euthanization, each zebrafish was weighed, photographed for measuring length using Image J. To determine fasting blood glucose levels, adult zebrafish were not fed in the morning and were collected to be euthanized using (MS-222) and clipped on the tail region for blood samples. The blood samples were then measured for using a CVS health glucose meter to determine blood glucose levels (mg/ml). For post-prandial glucose measurements, the fish were fed in the morning, and blood glucose was measured 1 hour post feeding. See **Supplementary Data S10** for raw data Prism file.

### Alamar blue assay

Zebrafish were maintained in 10-cm Petri dishes at 28°C from 0–3 dpf in E3 media ^52^. E3 media was replaced daily. At day 3 dpf, the larvae were rinsed with E3 and pipetted into a modified 96 well plate with 3 zebrafish larvae inside each well. Residual liquid from the transfer was pipetted out and 200 ul of alarm blue assay buffer was added to each well. The assay buffer is E3 supplemented with 1% (10×) Alamar Blue. Fluorescence of the plate was immediately read on a SpectraMax M5 multidetection reader (Molecular Devices, Sunnyvale, CA) with excitation at 530 nm and emission monitored at 590 nm. Measurements were taken at time intervals of 0,1,2,3,24, and 48 hours. Post-measurements, the plate was covered with aluminum foil and stored in a 28°C incubator until the next measurement time. Any wells that had fish die during the assay were excluded from analysis. Data were corrected by setting the average of the no drug wild-type control to 1.

Surface and Tinaja embryos were staged during collection due to variable time differences in fertilization of embryos. The Alamar blue assay was conducted at matching times of development between both genotypes to ensure that biases towards developmental stages was reduced and the same protocol was applied as described above.

For adult zebrafish, a published protocol was used ^31^. Briefly, individual fish were put in 50ml falcon tubes with 25-35ml of 0.02mg/ml Alamar blue solution made in system water and kept in 28°C incubator for up to 72 hours. During this time food was restricted as that can hinder with the assay. 100ul of Alamar Blue solution was sampled periodically to measure florescence at 590nm.

See **Supplementary Data S10** for raw data Prism file.

### Oil Red O staining

Zebrafish larvae at 3dpf and 5dpf were euthanized using MS-222 and stained with Oil Red O according to a published protocol ^53^. Post-euthanization, samples were collected and fixed overnight in 4% PFA at 2-8°C. The following day samples were washed using PBS/0.1% Tween-20 (PBSTw) and then incubated in 0.3% Oil Red O working solution for 15 minutes. The working solution was removed at 15 minutes and samples were washed in the following manner: PBSTw, 60% isopropanol, and a final wash of PBSTw.

The stain was quantified using ImageJ to measure the amount of stain present in the yolk region between both genotypes for comparisons.

See **Supplementary Data S10** for raw data Prism file.

### Feeding assay

Frozen blood worms from Hikari were cut into an appropriate size of 1.5 mm and 25 pieces were collected for each genotype and sex. Zebrafish were fed in the morning, and after a 1-hour interval, pieces were recounted to determine how many were consumed by each group. Blood worms were used in the assay for the ease of counting and measuring amounts of food consumed.

See **Supplementary Data S10** for raw data Prism file.

### Locomotion assays

Zebrafish at 3dpf were collected and transferred into a 96-well PCR plate and incubated for one day to acclimate. The following day, the plate was transferred to a DanioVision, which allowed tracking of movement for each genotype over a 3-hour incubation. The incubation was set for alternating 30-minute light/dark cycles to determine of there is a bias toward light/dark stimuli. In addition to this, Zebrafish at 3 dpf were transferred to a 24-well plate and video was recorded for 30 seconds. For tracking Astyanax, lack of contrast in the cavefish larvae prevented camera from detecting them. Hence, we waited till 8dpf when the fish could eat artemia. The artemia in their gut provided the necessary contrast allowing easy camera detection. *Zebra zoom* software was utilized to trace and determine the total distance travelled by each larva ^54^.

See **Supplementary Data S11 and S12** for raw data Prism files.

## Author Contributions

JK and NR conceived the project. JK, NR, and TG planned, and JK supervised the study. TG, TB, ET, KD, KW, GV, CP, and VT performed the experiments. MCR provided EchoMRI support. TG and JK wrote the manuscript. All authors read and approved the manuscript.

## Supporting information

Guzman_Supplementary Data

## Acknowledgements

We thank Comparative Medicine Core Facility and Sarah Ball for the zebrafish and cavefish husbandry at OMRF, CBDS Core for help with statistical analyses. We also thank the Aquatics Core Facility for the zebrafish husbandry at the Stowers Institute for Medical Research.

## Supplementary Figures

**Figure S1:**
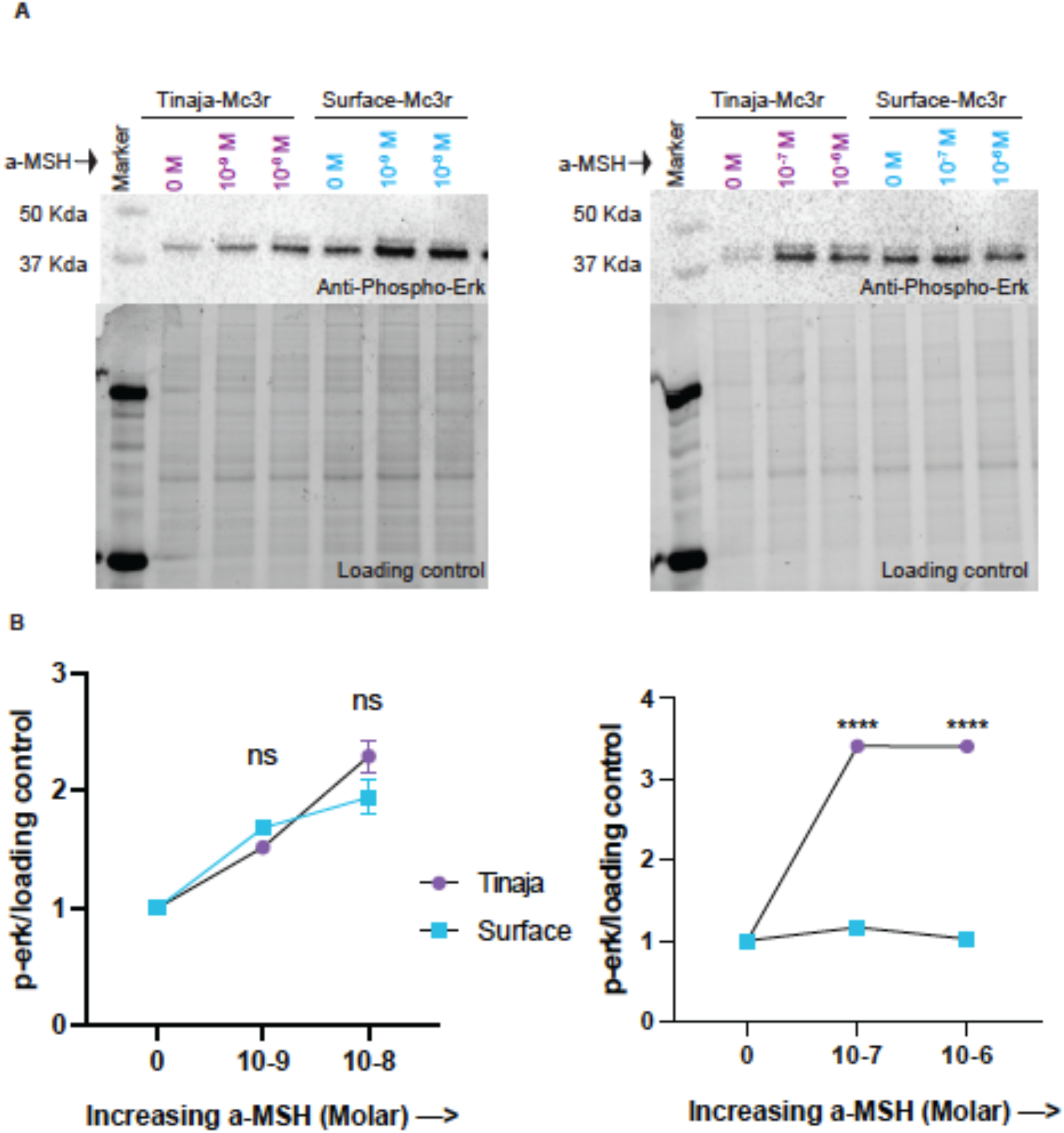
(A) p.ERK quantification: Upper panel shows a western blots for p-ERK on whole cell extracts of HEK 293T cells expressing either surface fish or Tmaja *mc3r* and stimulated with various concentrations of a-MSH ligand. Lower panel shows the image of stain-free gel to quantify protein loading amounts. (8) Densitometric quantification of *the* western blot signal normalized to the respective loading controls and the signal with OM a-MSH. The data were analyzed using Two way 1>1-JOVA followed by Sidak’s multiple comparison test. The data represents mean+_ SD with nsP>0.05, ****P<0.0001

**Figure S2:**
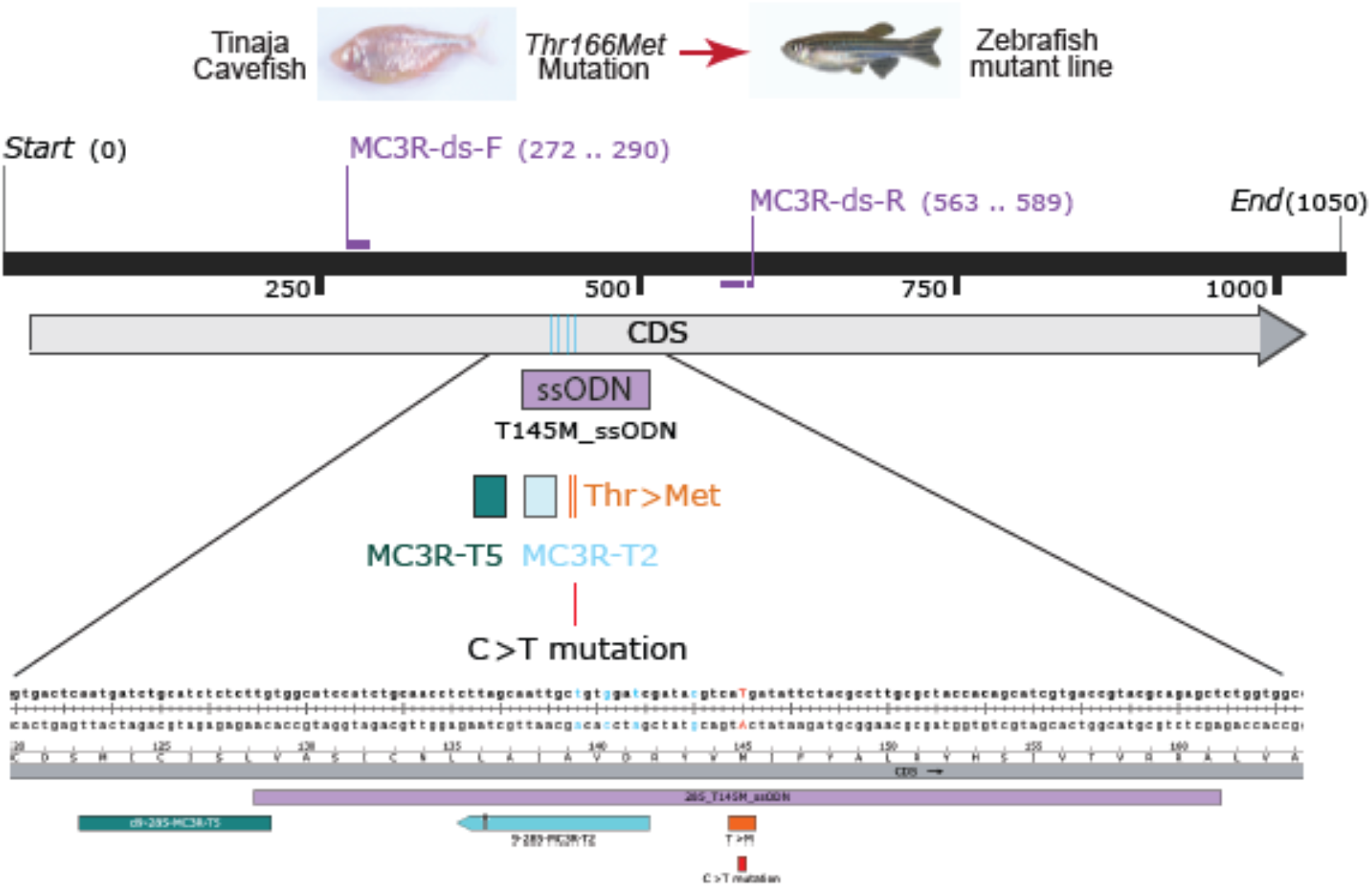
CRISPR scheme for generating the Thr166Met mutation. CDS is the coding sequence for mc3r. 9-285-MC3R-T2 and d9-285-MC3R-T5 are the gRNAs used. ssODN is the oligonucleotide used for homologous recombination. MC3R-ds-F and MC3R-ds-R are primers used for genotyping.

**Figure S3:**
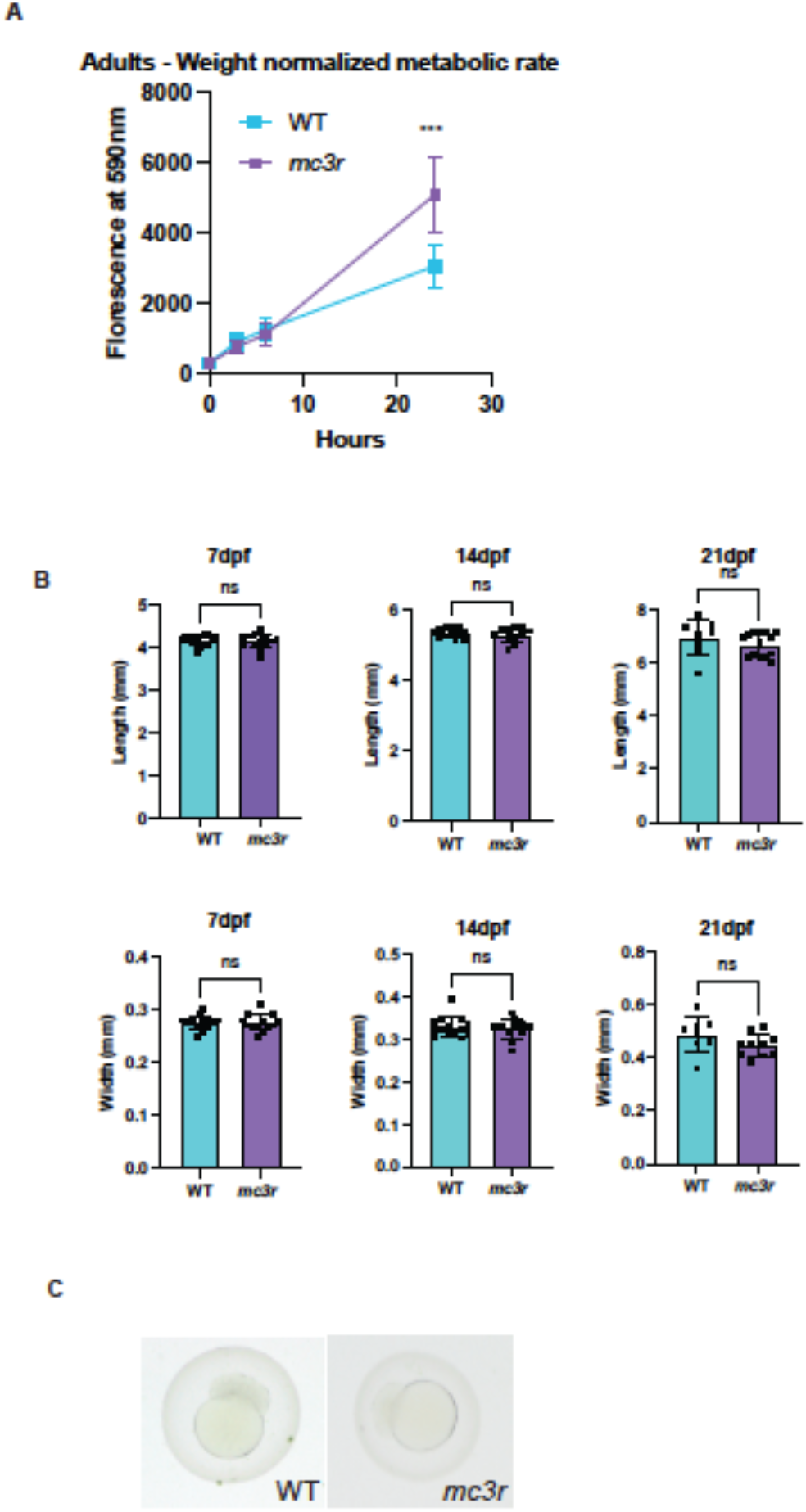
**(A)** Weight-normalize<l metabolic Rates of **WT** an<l mutant zebrafish measure<l using Alamar Blue assay. Age of fish - 15mpf. P value <0.001. **(B)** Length an<l HAA (Height at the Anterior of Anal Fin) for WT an<l mutants at S<lpf, 7dpf, 14<lpf, an<l 21Clpf. (C) Images of WT an<l mutant zebrafish embryos at 2hpf showing no significant differences in their development.

**Figure S4:**
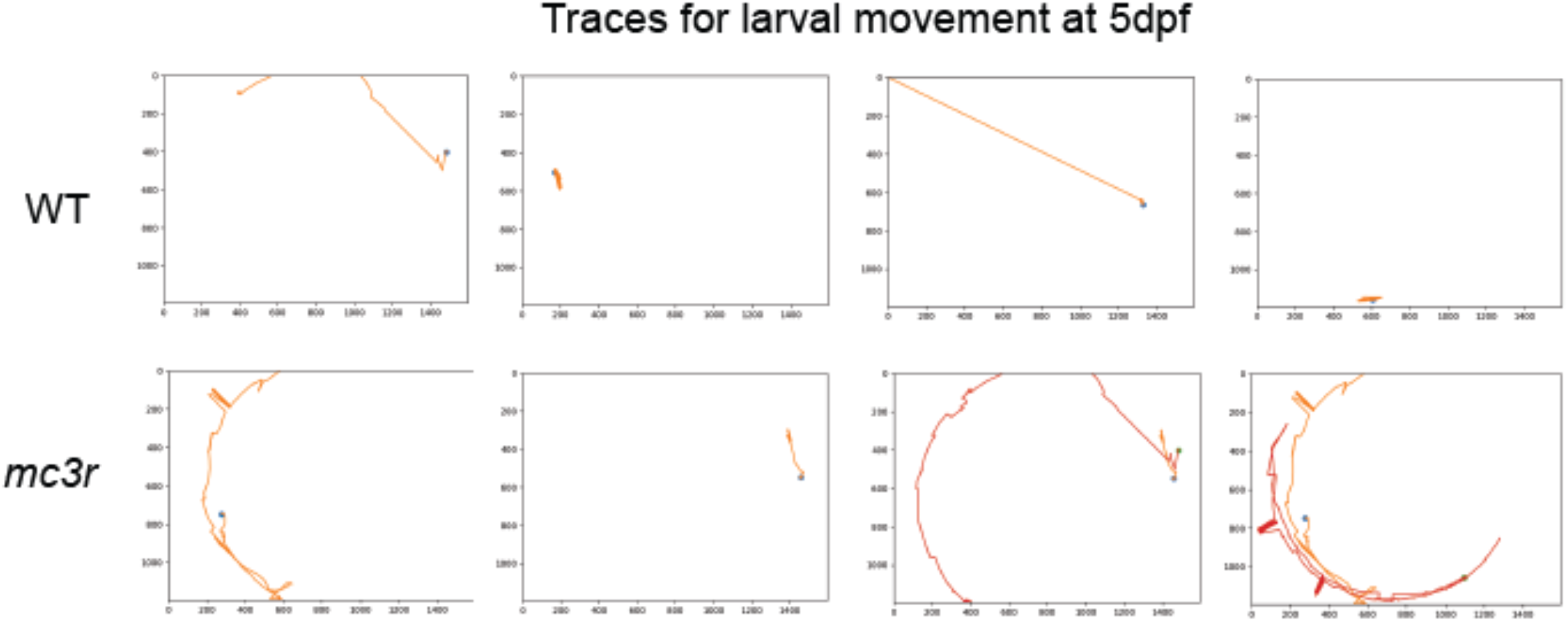
Movement traces of five WT and mutant zebrafish larvae at Sdpf. Images generated as a standard output from Zebra Zoom behavior tracking software.

