## Supplementary material for "A naturally evolved mutation in cavefish *mc3r* reshapes energy homeostasis across development in a nutrient-limited environment": Guzman_Supplementary Data: Supplementary_Data_S1_GPCR.pdf

**METHODS**

Starting with the complete set of named G-protein-coupled receptors (GPCRs) encoded by the *Astyanax mexicanus* surface reference assembly (AstMex3; GCF_023375975.1), defined as AstMex3 gene symbols matching the curated human G-protein-coupled-receptor superfamily (as defined by the HUGO Gene Nomenclature Committee gene group "G protein-coupled receptors"). They were matched by their ortholog symbol and teleost whole-genome-duplication ohnologs (e.g. mc5ra/mc5rb) were taken into account. After discarding unnamed (LOC) loci (n = 311 receptors), the remaining GPCRs were filtered to find those relevant to metabolism and/or the HPA (hypothalamic-pituitary-adrenal axis)/neuroendocrine system, the categories most relevant to the cavefish metabolic phenotype. Using the IUPHAR/BPS Guide to Pharmacology (GtoPdb, v2026.2), each receptor's human ortholog was mapped to its GtoPdb family, families were assigned to classes by their established physiology, and ranking of candidates of potential interest was restricted to these two classes.

Coding variants were drawn from the cave/surface cohort of 45 wild-caught individuals encompassing three cave populations (Tinaja, Pachón, Molino) and two surface populations (Río Mante, Río Rascón). They were re-aligned to AstMex3 and jointly genotyped (GATK), then filtered to high-quality (PASS) sites. Variants within the 54 receptors were annotated for coding consequence against a custom AstMex3 SnpEff database, using each gene's longest-protein isoform. For every variant, alternate-allele frequencies were computed for three groups: Tinaja, other-cave (Pachón + Molino), and surface (Mante + Rascón).

Only cave-specific missense variants were considered – amino-acid-changing substitutions present in at least one cave population and absent from surface (surface allele frequency = 0), so that every reported variant is a cave-specific coding change relative to the surface reference. The potential impact of each substitution was scored with the ESM-2 protein language model (650M-parameter masked language model, esm2_t33_650M_UR50D) on the native AstMex3 protein. The variant residue was masked and the log-likelihood ratio between the cave (alternate) and surface (reference) amino acid was read from a single forward pass. Its absolute value, |ESM-LLR|, is the magnitude of predicted perturbation.

Receptors were ranked by a composite score to weight: 1) Enrichment - the observed cave-versus-surface allele-frequency difference, max(AF_Tinaja, AF_other-cave) − AF_surface; and 2) |ESM-LLR|, the predicted functional perturbation. Both were standardized (z-scored) across the pool of all cave-specific missense variants and combined as a score = z(enrichment) + 0.5 × z(|F-imp|). For each gene, the variant chosen was the highest-scoring cave-specific missense variant. Enrichment is weighted more heavily than predicted impact because it is derived from directly observed population-genetic data, whereas |ESM-LLR| is a sequence-model prediction, which can only serve as a noisy proxy for the potential impact on function of a substitution (relative to surface). The goal was to rank mutations that were both cavefish-specific AND of predicted consequence.

**Table 1. Metabolic / neuroendocrine GPCRs ranked by cave-specific coding perturbation**

| Rank | score | Symbol | Ortholog | Pathway | # CSM | Highest-score variant | \|ESM-LLR\| | sign | Enrichment | AF Tinaja | AF other-cave | dxy max pct |
|---|---|---|---|---|---|---|---|---|---|---|---|---|
| **1** | 3.27 | hcrtr2 | HCRTR2 | metabolic | 4 | Glu295Lys | 8.37 | + | 0.955 | 0.955 | 0.583 | 80.2 |
| **2** | 2.38 | tacr3a | TACR3 | HPA_neuroendocrine | 2 | Val145Met | 3.47 | + | 0.955 | 0.955 | 0.000 | 93.2 |
| **3** | 2.34 | qrfprb | QRFPR | HPA_neuroendocrine | 5 | Arg417His | 2.48 | + | 1.000 | 1.000 | 0.000 | 96.1 |
| **4** | 2.29 | mc3r | MC3R | metabolic | 3 | Thr166Met | 3.76 | + | 0.909 | 0.909 | 0.000 | 92.8 |
| **5** | 2.11 | tacr1b | TACR1 | HPA_neuroendocrine | 2 | Leu180Ile | 2.01 | + | 0.955 | 0.955 | 0.529 | 89.9 |
| **6** | 2.08 | gcgrb | GCGR | metabolic | 3 | Ile300Phe | 1.08 | + | 1.000 | 1.000 | 0.000 | 97.5 |
| **7** | 2.06 | mchr1a | MCHR1 | metabolic | 4 | Glu354Ala | 0.98 | - | 1.000 | 1.000 | 0.647 | 90.6 |
| **8** | 1.93 | tacr2 | TACR2 | HPA_neuroendocrine | 2 | Leu98Ile | 0.26 | - | 1.000 | 1.000 | 0.700 | 71.1 |
| **9** | 1.92 | sstr1a | SSTR1 | HPA_neuroendocrine | 2 | Ala34Ser | 0.18 | - | 1.000 | 1.000 | 0.000 | 96.0 |
| **10** | 1.91 | cckbra | CCKBR | metabolic | 4 | Gly291Arg | 0.13 | + | 1.000 | 1.000 | 0.656 | 92.3 |
| **11** | 1.88 | mc4r | MC4R | metabolic | 2 | Val166Ile | 0.76 | + | 0.955 | 0.955 | 0.625 | 92.9 |
| **12** | 1.85 | npy4r | NPY4R | metabolic | 1 | Arg363His | 1.36 | + | 0.909 | 0.909 | 0.000 | 99.3 |
| **13** | 1.75 | gpr17 | GPR17 | metabolic | 2 | Ser152Gly | 3.53 | + | 0.750 | 0.667 | 0.750 | 88.1 |
| **14** | 1.71 | mc2r | MC2R | HPA_neuroendocrine | 1 | Lys169Asn | 0.61 | - | 0.909 | 0.909 | 0.714 | 86.8 |
| **15** | 1.68 | avpr1ab | AVPR1A | HPA_neuroendocrine | 1 | Leu77Met | 3.17 | + | 0.750 | 0.750 | 0.000 | 83.7 |
| **16** | 1.68 | cckar | CCKAR | metabolic | 2 | Glu404Lys | 1.99 | - | 0.818 | 0.818 | 0.000 | 74.9 |
| **17** | 1.46 | nmur1a | NMUR1 | HPA_neuroendocrine | 1 | Val152Met | 2.32 | + | 0.727 | 0.727 | 0.000 | 84.0 |
| **18** | 1.42 | galr1a | GALR1 | HPA_neuroendocrine | 7 | Val256Met | 4.95 | + | 0.562 | 0.562 | 0.000 | 91.2 |
| **19** | 1.02 | ghsra | GHSR | metabolic | 2 | His23Gln | 1.94 | - | 0.611 | 0.000 | 0.611 | 86.0 |
| **20** | 1.01 | avpr1aa | AVPR1A | HPA_neuroendocrine | 4 | Cys273Tyr | 9.70 | + | 0.154 | 0.000 | 0.154 | 93.3 |
| **21** | 0.89 | galr1b | GALR1 | HPA_neuroendocrine | 1 | Cys219Ser | 4.82 | - | 0.400 | 0.400 | 0.000 | 86.2 |
| **22** | 0.87 | trhrb | TRHR | HPA_neuroendocrine | 4 | Gln257Arg | 3.00 | + | 0.500 | 0.500 | 0.000 | 83.0 |
| **23** | 0.84 | npy2r | NPY2R | metabolic | 14 | Cys206Gly | 10.80 | + | 0.036 | 0.000 | 0.036 | 74.2 |
| **24** | 0.81 | galr2b | GALR2 | HPA_neuroendocrine | 10 | Val256Glu | 10.28 | + | 0.056 | 0.056 | 0.000 | 94.4 |
| **25** | 0.72 | npffr2a | NPFFR2 | HPA_neuroendocrine | 1 | Phe341Ser | 9.77 | + | 0.056 | 0.056 | 0.000 | 94.9 |
| **26** | 0.67 | sstr1b | SSTR1 | HPA_neuroendocrine | 1 | Glu245Lys | 9.98 | + | 0.028 | 0.000 | 0.028 | 100.0 |
| **27** | 0.67 | npy1r | NPY1R | metabolic | 3 | Arg271Cys | 8.31 | + | 0.125 | 0.000 | 0.125 | 97.0 |
| **28** | 0.54 | ghsrb | GHSR | metabolic | 11 | Thr60Pro | 8.67 | + | 0.062 | 0.062 | 0.000 | 72.7 |
| **29** | 0.42 | sstr2a | SSTR2 | HPA_neuroendocrine | 3 | Gln133Arg | 7.88 | + | 0.071 | 0.071 | 0.000 | 82.5 |
| **30** | 0.27 | aplnrb | APLNR | metabolic | 2 | Ile214Leu | 1.25 | - | 0.412 | 0.000 | 0.412 | 90.0 |
| **31** | 0.16 | oxtrb | OXTR | HPA_neuroendocrine | 9 | Lys228Met | 7.15 | + | 0.031 | 0.000 | 0.031 | 88.9 |
| **32** | 0.10 | oxtra | OXTR | HPA_neuroendocrine | 1 | Cys149Ser | 6.72 | + | 0.036 | 0.000 | 0.036 | 76.2 |
| **33** | 0.05 | gipr | GIPR | metabolic | 2 | His505Gln | 0.02 | + | 0.412 | 0.000 | 0.412 | 84.2 |
| **34** | 0.03 | cnr2 | CNR2 | metabolic | 1 | Gly21Glu | 0.03 | + | 0.406 | 0.000 | 0.406 | 89.9 |
| **35** | -0.09 | tacr1a | TACR1 | HPA_neuroendocrine | 1 | Ala167Ser | 3.86 | + | 0.143 | 0.143 | 0.000 | 94.4 |
| **36** | -0.21 | cnr1 | CNR1 | metabolic | 1 | Phe408Ser | 4.56 | + | 0.062 | 0.062 | 0.000 | 86.0 |
| **37** | -0.38 | npbwr2b | NPBWR2 | HPA_neuroendocrine | 3 | Ser149Phe | 4.17 | + | 0.031 | 0.000 | 0.031 | 95.9 |
| **38** | -0.41 | nmur1b | NMUR1 | HPA_neuroendocrine | 4 | Lys170Thr | 3.40 | + | 0.067 | 0.000 | 0.067 | 91.6 |

| | | | | | | | | | | | | |
|---|---|---|---|---|---|---|---|---|---|---|---|---|
| **39** | -0.44 | gcgra | GCGR | metabolic | 1 | Lys282Thr | 3.91 | + | 0.029 | 0.000 | 0.029 | 94.4 |
| **40** | -0.46 | mtnr1bb | MTNR1B | HPA_neuroendocrine | 2 | Val203Met | 2.80 | + | 0.088 | 0.000 | 0.088 | 89.5 |
| **41** | -0.65 | mtnr1aa | MTNR1A | HPA_neuroendocrine | 7 | Gly114Ala | 2.72 | - | 0.031 | 0.000 | 0.031 | 99.3 |
| **42** | -0.72 | gpr75 | GPR75 | metabolic | 1 | Thr17Lys | 2.34 | + | 0.031 | 0.000 | 0.031 | 89.5 |
| **43** | -0.72 | sstr3 | SSTR3 | HPA_neuroendocrine | 2 | Thr56Pro | 2.30 | + | 0.033 | 0.000 | 0.033 | 89.8 |
| **44** | -0.79 | ghrhra | GHRHR | HPA_neuroendocrine | 2 | Val205Ile | 1.43 | + | 0.062 | 0.062 | 0.000 | 93.9 |
| **45** | -0.87 | trhra | TRHR | HPA_neuroendocrine | 5 | Ser400Leu | 0.35 | + | 0.100 | 0.100 | 0.000 | 86.6 |
| **46** | -0.96 | mtnr1ab | MTNR1A | HPA_neuroendocrine | 1 | Leu315Arg | 0.75 | + | 0.045 | 0.045 | 0.033 | 93.4 |
| **47** | -1.00 | gpbar1 | GPBAR1 | metabolic | 1 | Ala278Val | 0.08 | - | 0.071 | 0.071 | 0.000 | 92.9 |
| **48** | -1.01 | mtnr1ba | MTNR1B | HPA_neuroendocrine | 2 | Pro16Ala | 0.73 | - | 0.033 | 0.000 | 0.033 | 83.8 |
| **49** | -1.05 | ghrhrb | GHRHR | HPA_neuroendocrine | 2 | Phe290Leu | 0.51 | - | 0.033 | 0.000 | 0.033 | 92.9 |
| **NA** | NA | aplnra | APLNR | metabolic | 0 | NA | NA | NA | NA | NA | NA | 74.8 |
| **NA** | NA | crhr1 | CRHR1 | HPA_neuroendocrine | 0 | NA | NA | NA | NA | NA | NA | 95.4 |
| **NA** | NA | kiss1ra | KISS1R | HPA_neuroendocrine | 0 | NA | NA | NA | NA | NA | NA | 93.5 |
| **NA** | NA | mchr1b | MCHR1 | metabolic | 0 | NA | NA | NA | NA | NA | NA | 93.2 |
| **NA** | NA | mchr2a | MCHR2 | metabolic | 0 | NA | NA | NA | NA | NA | NA | 91.0 |

**Legend:** Each row is one GPCR receptor; the variant shown is that receptor's single cave-specific missense substitution with the highest rank_score. **Score -** score = z(enrichment) + 0.5z(|F-imp|), where z=standardized z-score for all cave-specific missense (CSM) variants. **Symbol -** AstMex3 gene symbol (teleost ohnologs carry a/b suffixes). **Ortholog -** Human ortholog symbol (HGNC). **# CSM -** Number of cave-specific missense variants in the receptor. **|ESM-LLR| -** ESM-2, a protein language model, gives a masked-marginal log-likelihood ratio for each cave-specific missense (CSM) variant: the cave residue's likelihood versus the surface allele's at the masked position. |ESM-LLR| is the size of the divergence from the model's expectation. **Enrichment -** Cave-versus-surface allele-frequency difference of the CSM variant: max(AF_Tinaja, AF_other-cave) − AF_surface. Equals the maximum cave allele frequency, since surface frequency = 0 for all rows. **AF Tinaja -** Alternate-allele frequency of the CSM variant in the Tinaja population. **AF other-cave -** Alternate-allele frequency in the combined Pachón + Molino populations. **dxy max pct -** Maximum windowed d_xy percentile across the gene over the six cave-surface population pairs (genome-wide divergence context).
