## Supplementary material for "A naturally evolved mutation in cavefish *mc3r* reshapes energy homeostasis across development in a nutrient-limited environment": Guzman_Supplementary Data: Supplementary_Data_S2_mc3r_BlastP.pdf

>Human_MC3R

MSIQKTYLEGDFVFPVSSSSFLRTLLEPQLGSALLTAMNASCCLPSVQPTLPNGSEHLQAPFFSNQS
SSAFCEQVFIKPEVFLSLGIVSLLENILVILAVVRNGNLHSPMYFFLCSLAVADMLVSVSNALETIMIAI
VHSDYLTFEDQFIQHMDNIFDSMICISLVASICNLLAIAVDRYVTIFYALRYHSIMTVRKALTLIVAIWV
CCGVCGVVFIVYSESKMVIVCLITMFFAMMLLMGTLYVHMFLFARLHVKRIAALPPADGVAPQQHS
CMKGAVTITILLGVFIFCWAPFFLHLVLIITCPTNPYCICYTAHFNTYLVLIMCNSVIDPLIYAFRSLELR
NTFREILCGCNGMNLG*

>Mouse_MC3R

MNSSCCLSSVSPMLPNLSEHPAAPPASNRSGSGFCEQVFIKPEVFLALGIVSLMENILVILAVVRNG
NLHSPMYFFLCSLAAADMLVSLSNSLETIMIAVINSDSLTLEDQFIQHMDNIFDSMICISLVASICNLL
AIAIDRYVTIFYALRYHSIMTVRKALTLIGVIWVCCGICGVMFIIYSESKMVIVCLITMFFAMVLLMGTLYI
HMFLFARLHVQRIAVLPPAGVVAPQQHSCMKGAVTITILLGVFIFCWAPFFLHLVLIITCPTNPYCICY
TAHFNTYLVLIMCNSVIDPLIYAFRSLELRNTFKEILCGCNSMNLG*

>Zebrafish_MC3R

MNDSHLQFLKGQKSVNSTSLPPNGSLADSPAGTLCEQVQIQAEVFLTLGIVSLLENILVISAVVKNKN
LHSPMYFFLCSLAAADMLVSVSNSLETIVIAVLNSRLVVASDQFVRLMDNVCDSMICISLVASICNLL
AIAVDRYVTIFYALRYHSIVTVRRALVAIAVIWLVCVVCGIVFIVYSESKTVIVCLITMFFAMLVLMATLYV
HMFLLARLHVQRIAALPPAAPGAGNPAPRQRSCMKGAVTISILLGVFVCCWAPFFLHLILLVSCPHH
PLCLCYMSHFTTYLVLIMCNSVIDPLIYACRSLEMRKTFKEILCCFGCQPAL*

>Surface_Astyanax

MNGSHHPVLLLLLQESSNTNLTEESEAAAASEPSNGTAPALVSNSSTAPPPAGALCQQVQIQAEVFL
ALGIVSLLENILVISAVAKNKNLHSPMYFFLCSLAAADMLVSVSNSLETIVIAVLRSHLLELSDYFVRLM
DNVFDSMICISLVASICNLLAIAVDRYVTIFYALRYHSIVTVRRALNAIAAIWLVCIVCGIVFIVYSESKTV
IICLITMFFAMLALMATLYVHMFLLARLHVQRIAALPPAASAAGNRAPRQQHTCMKGAVTITILLGVFV
CCWAPFFLHLILLVACPQHPLCLCYMSHFTTYLVLIMCNSVIDPIIYAFRSLEMRKTFKEILCCFGESC
QAPLCERERERERERDTETERSRERQHEREKESETE*

>Pachòn_Astyanax

MNGSHHPVLLLLVQESSNTNLTEESEAAAASEPSNGTAPALVSNSSTAPPPAGALCQQVQIQAEVF
LALGIVSLLENILVISAVAKNKNLHSPMYFFLCSLAAADMLVSVSNSLETIMIAVLRSHLLELNDYFVR
LMDNVFDSMICISLVASICNLLAIAVDRYVTIFYALRYHSIVTVRRALNAIAAIWLVCIVCGIVFIVYSES
KTVIICLITMFFAMLALMATLYVHMFLLARLHVQRIAALPPAASAAGNRAPRQQHTCMKGAVTITILLG
VFVCCWAPFFLHLILLVACPQHPLCLCYMSHFTTYLVLIMCNSVIDPIIYAFRSLEMRKTFKEILCCFG
ESCQAPLCERERERERERDTETERSRERQHEREKESETE*

>Tinaja_Astyanax

MNGSHHPVLLLLLQESSNTNLTEESEAAAASEPSNGTAPALVSNSSTAPPPAGALCQQVQIQAEVFL
ALGIVSLLENILVISAVAKNKNLHSPMYFFLCSLAAADMLVSVSNSLETIVIAVLRSHLLELSDYFVRLM
DNVFDSMICISLVASICNLLAVAVDRYVMIFYALRYHSIVTVRRALNAIAAIWLVCIVCGIVFIVYSESKT
VIICLITMFFAMLALMATLYVHMFLLARLHVQRIAALPPAASAAGNPAPRQQHTCMKGAVTITILLGVF
VCCWAPFFLHLILLVACPQHPLCLCYMSHFTTYLVLIMCNSVIDPIIYAFRSLEMRKTFKEILCCFGES
GQAPLCERERERERERDTETERSRERQREREKESETE*

```
CLUSTAL O(1.2.4) multiple sequence alignment


Human_mc3r          ---------------------------LPSVQPTLPNGSEHLQAPFFSNQSSSAFCEQVF        33
Mouse_mc3r          MNSS---------------------CCLSSVSPMLPNLSEHPAAPPASNRSGSGFCEQVF        39
Zebrafish_mc3r      MNDSHLQ------------FLK---------GQKSVNSTSLPPNGSLADSPAGTLCEQVQ        39
Surface_mc3r        MNGSHHPVLLLLLQESSNTNLTEESEAAAASEPSNGTAPALVSNSSTAPPPAGALCQQVQ        60
Pachon_mc3r         MNGSHHPVLLLLVQESSNTNLTEESEAAAASEPSNGTAPALVSNSSTAPPPAGALCQQVQ        60
Tinaja_mc3r         MNGSHHPVLLLLLQESSNTNLTEESEAAAASEPSNGTAPALVSNSSTAPPPAGALCQQVQ        60
                                                    .          :   .. :*:** 

Human_mc3r          IKPEVFLSLGIVSLLENILVILAVVRNGNLHSPMYFFLCSLAVADMLVSVSNALETIMIA        93
Mouse_mc3r          IKPEVFLALGIVSLMENILVILAVVRNGNLHSPMYFFLCSLAAADMLVSLSNSLETIMIA        99
Zebrafish_mc3r      IQAEVFLTLGIVSLLENILVISAVVKNKNLHSPMYFFLCSLAAADMLVSVSNSLETIVIA        99
Surface_mc3r        IQAEVFLALGIVSLLENILVISAVAKNKNLHSPMYFFLCSLAAADMLVSVSNSLETIVIA        120
Pachon_mc3r         IQAEVFLALGIVSLLENILVISAVAKNKNLHSPMYFFLCSLAAADMLVSVSNSLETIMIA        120
Tinaja_mc3r         IQAEVFLALGIVSLLENILVISAVAKNKNLHSPMYFFLCSLAAADMLVSVSNSLETIVIA        120
                    *: ****:******:****** **.:* **************.******:**:****:**

Human_mc3r          IVHSDYLTFEDQFIQHMDNIFDSMICISLVASICNLLAIAVDRYVTIFYALRYHSIMTVR        153
Mouse_mc3r          VINSDSLTLEDQFIQHMDNIFDSMICISLVASICNLLAIAIDRYVTIFYALRYHSIMTVR        159
Zebrafish_mc3r      VLNSRLVVASDQFVRLMDNVCDSMICISLVASICNLLAIAVDRYVTIFYALRYHSIVTVR        159
Surface_mc3r        VLRSHLLELSDYFVRLMDNVFDSMICISLVASICNLLAIAVDRYVTIFYALRYHSIVTVR        180
Pachon_mc3r         VLRSHLLELNDYFVRLMDNVFDSMICISLVASICNLLAIAVDRYVTIFYALRYHSIVTVR        180
Tinaja_mc3r         VLRSHLLELSDYFVRLMDNVFDSMICISLVASICNLLAVAVDRYVMIFYALRYHSIVTVR        180
                    ::.*  :  .* *:: ***: *****************:*:**** **********:***

Human_mc3r          KALTLIVAIWVCCGVCGVVFIVYSESKMVIVCLITMFFAMMLLMGTLYVHMFLFARLHVK        213
Mouse_mc3r          KALTLIGVIWVCCGICGVMFIIYSESKMVIVCLITMFFAMVLLMGTLYIHMFLFARLHVQ        219
Zebrafish_mc3r      RALVAIAVIWLVCVVCGIVFIVYSESKTVIVCLITMFFAMLVLMATLYVHMFLLARLHVQ        219
Surface_mc3r        RALNAIAAIWLVCIVCGIVFIVYSESKTVIICLITMFFAMLALMATLYVHMFLLARLHVQ        240
Pachon_mc3r         RALNAIAAIWLVCIVCGIVFIVYSESKTVIICLITMFFAMLALMATLYVHMFLLARLHVQ        240
Tinaja_mc3r         RALNAIAAIWLVCIVCGIVFIVYSESKTVIICLITMFFAMLALMATLYVHMFLLARLHVQ        240
                    :**  * .**: * :**::**:***** **:*********: **.***:****:*****:

Human_mc3r          RIAALPPADGV-----APQQHSCMKGAVTITILLGVFIFCWAPFFLHLVLIITCPTNPYC        268
Mouse_mc3r          RIAVLPPAGVV-----APQQHSCMKGAVTITILLGVFIFCWAPFFLHLVLIITCPTNPYC        274
Zebrafish_mc3r      RIAALPPAAPGAGNPAPRQ-RSCMKGAVTISILLGVFVCCWAPFFLHLILLVSCPHHPLC        278
Surface_mc3r        RIAALPPAASAAGNRAPRQQHTCMKGAVTITILLGVFVCCWAPFFLHLILLVACPQHPLC        300
Pachon_mc3r         RIAALPPAASAAGNRAPRQQHTCMKGAVTITILLGVFVCCWAPFFLHLILLVACPQHPLC        300
Tinaja_mc3r         RIAALPPAASAAGNPAPRQQHTCMKGAVTITILLGVFVCCWAPFFLHLILLVACPQHPLC        300
                    ***.****          * ::********:******: *********:*:::** :* *

Human_mc3r          ICYTAHFNTYLVLIMCNSVIDPLIYAFRSLELRNTFREILCGCNGMNLG*----------        317
Mouse_mc3r          ICYTAHFNTYLVLIMCNSVIDPLIYAFRSLELRNTFKEILCGCNSMNLG*----------        323
Zebrafish_mc3r      LCYMSHFTTYLVLIMCNSVIDPLIYACRSLEMRKTFKEILCCFGCQPAL*----------        327
Surface_mc3r        LCYMSHFTTYLVLIMCNSVIDPIIYAFRSLEMRKTFKEILCCFGESCQAPLCERERERER        360
Pachon_mc3r         LCYMSHFTTYLVLIMCNSVIDPIIYAFRSLEMRKTFKEILCCFGESCQAPLCERERERER        360
Tinaja_mc3r         LCYMSHFTTYLVLIMCNSVIDPIIYAFRSLEMRKTFKEILCCFGESGQAPLCERERERER        360
                    :** :**.**************:*** ****:*:**:****  .

Human_mc3r          ------------------------    317
Mouse_mc3r          ------------------------    323
Zebrafish_mc3r      ------------------------    327
Surface_mc3r        ERDTETERSRERQHEREKESETE*    383
```

```
CLUSTAL O(1.2.4) multiple sequence alignment


Surface_mc3r      MNGSHHPVLLLLLQESSNTNLTEESEAAAASEPSNGTAPALVSNSSTAPPPAGALCQQVQ  60
Pachon_mc3r       MNGSHHPVLLLLVQESSNTNLTEESEAAAASEPSNGTAPALVSNSSTAPPPAGALCQQVQ  60
Tinaja_mc3r       MNGSHHPVLLLLLQESSNTNLTEESEAAAASEPSNGTAPALVSNSSTAPPPAGALCQQVQ  60
                  ************:***********************************************

Surface_mc3r      IQAEVFLALGIVSLLENILVISAVAKNKNLHSPMYFFLCSLAAADMLVSVSNSLETIVIA  120
Pachon_mc3r       IQAEVFLALGIVSLLENILVISAVAKNKNLHSPMYFFLCSLAAADMLVSVSNSLETIMIA  120
Tinaja_mc3r       IQAEVFLALGIVSLLENILVISAVAKNKNLHSPMYFFLCSLAAADMLVSVSNSLETIVIA  120
                  *********************************************************:**

Surface_mc3r      VLRSHLLELSDYFVRLMDNVFDSMICISLVASICNLLAIAVDRYVTIFYALRYHSIVTVR  180
Pachon_mc3r       VLRSHLLELNDYFVRLMDNVFDSMICISLVASICNLLAIAVDRYVTIFYALRYHSIVTVR  180
Tinaja_mc3r       VLRSHLLELSDYFVRLMDNVFDSMICISLVASICNLLAVAVDRYVMIFYALRYHSIVTVR  180
                  *********.****************************:****** **************

Surface_mc3r      RALNAIAAIWLVCIVCGIVFIVYSESKTVIICLITMFFAMLALMATLYVHMFLLARLHVQ  240
Pachon_mc3r       RALNAIAAIWLVCIVCGIVFIVYSESKTVIICLITMFFAMLALMATLYVHMFLLARLHVQ  240
Tinaja_mc3r       RALNAIAAIWLVCIVCGIVFIVYSESKTVIICLITMFFAMLALMATLYVHMFLLARLHVQ  240
                  ************************************************************

Surface_mc3r      RIAALPPAASAAGNRAPRQQHTCMKGAVTITILLGVFVCCWAPFFLHLILLVACPQHPLC  300
Pachon_mc3r       RIAALPPAASAAGNRAPRQQHTCMKGAVTITILLGVFVCCWAPFFLHLILLVACPQHPLC  300
Tinaja_mc3r       RIAALPPAASAAGNPAPRQQHTCMKGAVTITILLGVFVCCWAPFFLHLILLVACPQHPLC  300
                  ************** *********************************************

Surface_mc3r      LCYMSHFTTYLVLIMCNSVIDPIIYAFRSLEMRKTFKEILCCFGESCQAPLCERERERER  360
Pachon_mc3r       LCYMSHFTTYLVLIMCNSVIDPIIYAFRSLEMRKTFKEILCCFGESCQAPLCERERERER  360
Tinaja_mc3r       LCYMSHFTTYLVLIMCNSVIDPIIYAFRSLEMRKTFKEILCCFGESGQAPLCERERERER  360
                  ********************************************** *************

Surface_mc3r      ERDTETERSRERQHEREKESETE*      383
Pachon_mc3r       ERDTETERSRERQHEREKESETE*      383
Tinaja_mc3r       ERDTETERSRERQREREKESETE*      383
                  *************:**********
```

If you use this service, please consider citing the following publication: The EMBL-EBI Job Dispatcher sequence analysis tools framework in 2024. More information about this
