## Supplementary material for "A naturally evolved mutation in cavefish *mc3r* reshapes energy homeostasis across development in a nutrient-limited environment": Guzman_Supplementary Data: Supplementary_Data_S3_pcDNA-FRT-NFLAG-MC3R.pdf

**NEBuilder Assembly Tool v1.12.17**

Timestamp: 8/8/2018 @ 17:25

**User Selected Settings**

| | |
|---|---|
| **Product Version** | E5520 - NEBuilder High-Fidelity DNA Assembly Cloning Kit |
| **No. of Fragments** | 2-3 fragments (including vector) |
| **Construct Length** | less than 10 Kb |
| **Min. Overlap** | 20 bp |
| **Allow linear assemblies** | No |
| **PCR Polymerase** | Q5 High-Fidelity DNA Polymerase |
| **PCR Primer Conc.** | 500 nM |
| **Min. Primer Length** | 18 nt |

**Fragment Arrangement**

**Vector Digestion**

Vector backbone digested with PacI and PmeI

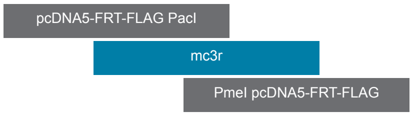


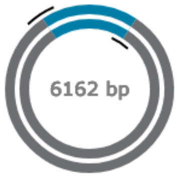


**Required Primers**

| Overlaps | Oligo (Uppercase = gene-specific primer) | Anneals | F/R | 3' Tm | 3' Ta * | 6-Frame |
|---|---|---|---|---|---|---|
| pcDNA5-FRT-FLAG | gacgatgatgacaagttaattaaaATGAACGGATCTCACCAC | mc3r | Fwd | 61.3°C | **62.3°C** | view |
| pcDNA5-FRT-FLAG | cgaggctgatcagcgggtttaaacTCACTCCGTCTCGCTCTC | mc3r | Rev | 66.2°C | **62.3°C** | view |

* 3' Ta (recommended annealing temperature for PCR) is calculated for the gene-specific portion of the primer for use with the selected PCR polymerase.

**Notes**

- Any element of a construction that includes the 5' overhang of a restriction site will be altered upon assembly. For example, the essential cysteine codon at the N-terminus of the intein segment of IMPACT vectors is present in the 5' overhang of the SapI site in those vectors. The bases removed in the assembly reaction can be added back by including them in the PCR primers for the corresponding insert.

**Assembled Sequence**

```
TCATAGCTCACGCTGTAGGTATCTCAGTTCGGTGTAGGTCGTTCGCTCCAAGCTGGGCTGTG
TGCACGAACCCCCCGTTCAGCCCGACCGCTGCGCCTTATCCGGTAACTATCGTCTTGAGTCC
AACCCGGTAAGACACGACTTATCGCCACTGGCAGCAGCCACTGGTAACAGGATTAGCAGAGC
GAGGTATGTAGGCGGTGCTACAGAGTTCTTGAAGTGGTGGCCTAACTACGGCTACACTAGAA
GGACAGTATTTGGTATCTGCGCTCTGCTGAAGCCAGTTACCTTCGGAAAAAGAGTTGGTAGC
TCTTGATCCGGCAAACAAACCACCGCTGGTAGCGGTGGTTTTTTTGTTTGCAAGCAGCAGAT
TACGCGCAGAAAAAAAGGATCTCAAGAAGATCCTTTGATCTTTTCTACGGGGTCTGACGCTC
AGTGGAACGAAAACTCACGTTAAGGGATTTTGGTCATGAGATTATCAAAAAGGATCTTCACC
TAGATCCTTTTAAATTAAAAATGAAGTTTTAAATCAATCTAAAGTATATATGAGTAAACTTG
GTCTGACAGTTACCAATGCTTAATCAGTGAGGCACCTATCTCAGCGATCTGTCTATTTCGTT
CATCCATAGTTGCCTGACTCCCCGTCGTGTAGATAACTACGATACGGGAGGGCTTACCATCT
GGCCCCAGTGCTGCAATGATACCGCGAGACCCACGCTCACCGGCTCCAGATTTATCAGCAAT
AAACCAGCCAGCCGGAAGGGCCGAGCGCAGAAGTGGTCCTGCAACTTTATCCGCCTCCATCC
AGTCTATTAATTGTTGCCGGGAAGCTAGAGTAAGTAGTTCGCCAGTTAATAGTTTGCGCAAC
GTTGTTGCCATTGCTACAGGCATCGTGGTGTCACGCTCGTCGTTTGGTATGGCTTCATTCAG
CTCCGGTTCCCAACGATCAAGGCGAGTTACATGATCCCCCATGTTGTGCAAAAAAGCGGTTA
GCTCCTTCGGTCCTCCGATCGTTGTCAGAAGTAAGTTGGCCGCAGTGTTATCACTCATGGTT
ATGGCAGCACTGCATAATTCTCTTACTGTCATGCCATCCGTAAGATGCTTTTCTGTGACTGG
TGAGTACTCAACCAAGTCATTCTGAGAATAGTGTATGCGGCGACCGAGTTGCTCTTGCCCGG
CGTCAATACGGGATAATACCGCGCCACATAGCAGAACTTTAAAAGTGCTCATCATTGGAAAA
CGTTCTTCGGGGCGAAAACTCTCAAGGATCTTACCGCTGTTGAGATCCAGTTCGATGTAACC
CACTCGTGCACCCAACTGATCTTCAGCATCTTTTACTTTCACCAGCGTTTCTGGGTGAGCAA
AAACAGGAAGGCAAAATGCCGCAAAAAAGGGAATAAGGGCGACACGGAAATGTTGAATACTC
ATACTCTTCCTTTTTCAATATTATTGAAGCATTTATCAGGGTTATTGTCTCATGAGCGGATA
CATATTTGAATGTATTTAGAAAAATAAACAAATAGGGGTTCCGCGCACATTTCCCCGAAAAG
TGCCACCTGACGTCGACGGATCGGGAGATCTCCCGATCCCCTATGGTGCACTCTCAGTACAA
TCTGCTCTGATGCCGCATAGTTAAGCCAGTATCTGCTCCCTGCTTGTGTGTTGGAGGTCGCT
GAGTAGTGCGCGAGCAAAATTTAAGCTACAACAAGGCAAGGCTTGACCGACAATTGCATGAA
GAATCTGCTTAGGGTTAGGCGTTTTGCGCTGCTTCGCGATGTACGGGCCAGATATACGCGTT
GACATTGATTATTGACTAGTTATTAATAGTAATCAATTACGGGGTCATTAGTTCATAGCCCA
TATATGGAGTTCCGCGTTACATAACTTACGGTAAATGGCCCGCCTGGCTGACCGCCCAACGA
CCCCCGCCCATTGACGTCAATAATGACGTATGTTCCCATAGTAACGCCAATAGGGACTTTCC
ATTGACGTCAATGGGTGGAGTATTTACGGTAAACTGCCCACTTGGCAGTACATCAAGTGTAT
CATATGCCAAGTACGCCCCCTATTGACGTCAATGACGGTAAATGGCCCGCCTGGCATTATGC
CCAGTACATGACCTTATGGGACTTTCCTACTTGGCAGTACATCTACGTATTAGTCATCGCTA
TTACCATGGTGATGCGGTTTTGGCAGTACATCAATGGGCGTGGATAGCGGTTTGACTCACGG
GGATTTCCAAGTCTCCACCCCATTGACGTCAATGGGAGTTTGTTTTGGCACCAAAATCAACG
GGACTTTCCAAAATGTCGTAACAACTCCGCCCCATTGACGCAAATGGGCGGTAGGCGTGTAC
GGTGGGAGGTCTATATAAGCAGAGCTCTCTGGCTAACTAGAGAACCCACTGCTTACTGGCTT
ATCGAAATTAATACGACTCACTATAGGGAGACCCAAGCTGGCTAGCATCATGCCAGACTACA
AGGACGATGATGACAAGTTAATTAAAATGAACGGATCTCACCACCCGGTGCTCCTCCTCCTC
CTGCAGGAAAGCTCCAACACCAACCTCACCGAAGAATCAGAAGCAGCAGCAGCTTCAGAACC
GAGCAATGGAACTGCACCAGCACTGGTTTCCAACAGCAGTACCGCCCCTCCACCAGCGGGGG
```

```
CGCTGTGCCAGCAGGTTCAGATCCAGGCGGAGGTGTTCCTCGCGCTCGGCATCGTCAGCCTG
CTGGAGAACATCCTGGTCATCTCGGCGGTGGCGAAGAACAAGAACCTCCACTCGCCCATGTA
CTTCTTTCTCTGTAGCTTGGCGGCGGCGGACATGCTGGTGAGTGTGTCCAACTCTCTGGAGA
CCATCGTGATCGCGGTTTTGAGGAGCCACCTGCTGGAGCTGAGCGATTATTTCGTGAGGCTG
ATGGACAATGTCTTCGACTCCATGATCTGCATCTCGCTGGTGGCGTCCATTTGCAACCTGCT
GGCGATCGCTGTCGATCGATACGTGACGATATTCTACGCTCTGCGTTACCACAGCATCGTTA
CGGTGCGGAGGGCGTTGAACGCCATTGCTGCCATCTGGCTGGTGTGCATCGTTTGCGGGATC
GTCTTCATCGTGTACTCCGAGAGTAAGACCGTCATCATCTGTCTGATCACCATGTTCTTCGC
CATGCTGGCGCTGATGGCCACCCTGTACGTGCACATGTTCCTCCTGGCCAGGCTCCACGTCC
AGCGCATCGCCGCGCTTCCCCCCGCGGCCAGCGCTGCAGGAAACCGTGCGCCGCGGCAGCAG
CACACCTGCATGAAGGGGGCGGTGACCATTACCATTCTCCTGGGCGTGTTCGTCTGCTGCTG
GGCGCCGTTCTTCCTGCACCTCATCCTGCTGGTGGCCTGCCCCCAGCACCCGCTCTGCCTGT
GCTACATGTCCCACTTCACCACCTACCTGGTGCTCATCATGTGCAACTCCGTCATCGACCCC
ATCATCTACGCCTTCCGCAGCCTGGAGATGAGGAAGACCTTCAAGGAAATCCTGTGCTGCTT
CGGCGAGAGCTGCCAAGCGCCGCTCTGCGAGAGAGAGCGCGAGAGAGAGCGCGAGAGAGACA
CAGAAACCGAGCGGTCGAGAGAGAGGCAGCACGAGAGGGAGAAGGAGAGCGAGACGGAGTGA
GTTTAAACCCGCTGATCAGCCTCGACTGTGCCTTCTAGTTGCCAGCCATCTGTTGTTTGCCC
CTCCCCCGTGCCTTCCTTGACCCTGGAAGGTGCCACTCCCACTGTCCTTTCCTAATAAAATG
AGGAAATTGCATCGCATTGTCTGAGTAGGTGTCATTCTATTCTGGGGGGTGGGGTGGGGCAG
GACAGCAAGGGGGAGGATTGGGAAGACAATAGCAGGCATGCTGGGGATGCGGTGGGCTCTAT
GGCTTCTGAGGCGGAAAGAACCAGCTGGGGCTCTAGGGGGTATCCCCACGCGCCCTGTAGCG
GCGCATTAAGCGCGGCGGGTGTGGTGGTTACGCGCAGCGTGACCGCTACACTTGCCAGCGCC
CTAGCGCCCGCTCCTTTCGCTTTCTTCCCTTCCTTTCTCGCCACGTTCGCCGGCTTTCCCCG
TCAAGCTCTAAATCGGGGGCTCCCTTTAGGGTTCCGATTTAGTGCTTTACGGCACCTCGACC
CCAAAAAACTTGATTAGGGTGATGGTTCACGTACCTAGAAGTTCCTATTCCGAAGTTCCTAT
TCTCTAGAAAGTATAGGAACTTCCTTGGCCAAAAAGCCTGAACTCACCGCGACGTCTGTCGA
GAAGTTTCTGATCGAAAAGTTCGACAGCGTCTCCGACCTGATGCAGCTCTCGGAGGGCGAAG
AATCTCGTGCTTTCAGCTTCGATGTAGGAGGGCGTGGATATGTCCTGCGGGTAAATAGCTGC
GCCGATGGTTTCTACAAAGATCGTTATGTTTATCGGCACTTTGCATCGGCCGCGCTCCCGAT
TCCGGAAGTGCTTGACATTGGGGAATTCAGCGAGAGCCTGACCTATTGCATCTCCCGCCGTG
CACAGGGTGTCACGTTGCAAGACCTGCCTGAAACCGAACTGCCCGCTGTTCTGCAGCCGGTC
GCGGAGGCCATGGATGCGATCGCTGCGGCCGATCTTAGCCAGACGAGCGGGTTCGGCCCATT
CGGACCGCAAGGAATCGGTCAATACACTACATGGCGTGATTTCATATGCGCGATTGCTGATC
CCCATGTGTATCACTGGCAAACTGTGATGGACGACACCGTCAGTGCGTCCGTCGCGCAGGCT
CTCGATGAGCTGATGCTTTGGGCCGAGGACTGCCCCGAAGTCCGGCACCTCGTGCACGCGGA
TTTCGGCTCCAACAATGTCCTGACGGACAATGGCCGCATAACAGCGGTCATTGACTGGAGCG
AGGCGATGTTCGGGGATTCCCAATACGAGGTCGCCAACATCTTCTTCTGGAGGCCGTGGTTG
GCTTGTATGGAGCAGCAGACGCGCTACTTCGAGCGGAGGCATCCGGAGCTTGCAGGATCGCC
GCGGCTCCGGGCGTATATGCTCCGCATTGGTCTTGACCAACTCTATCAGAGCTTGGTTGACG
GCAATTTCGATGATGCAGCTTGGGCGCAGGGTCGATGCGACGCAATCGTCCGATCCGGAGCC
GGGACTGTCGGGCGTACACAAATCGCCCGCAGAAGCGCGGCCGTCTGGACCGATGGCTGTGT
AGAAGTACTCGCCGATAGTGGAAACCGACGCCCCAGCACTCGTCCGAGGGCAAAGGAATAGC
ACGTACTACGAGATTTCGATTCCACCGCCGCCTTCTATGAAAGGTTGGGCTTCGGAATCGTT
TTCCGGGACGCCGGCTGGATGATCCTCCAGCGCGGGGATCTCATGCTGGAGTTCTTCGCCCA
CCCCAACTTGTTTATTGCAGCTTATAATGGTTACAAATAAAGCAATAGCATCACAAATTTCA
CAAATAAAGCATTTTTTTCACTGCATTCTAGTTGTGGTTTGTCCAAACTCATCAATGTATCT
TATCATGTCTGTATACCGTCGACCTCTAGCTAGAGCTTGGCGTAATCATGGTCATAGCTGTT
TCCTGTGTGAAATTGTTATCCGCTCACAATTCCACACAACATACGAGCCGGAAGCATAAAGT
GTAAAGCCTGGGGTGCCTAATGAGTGAGCTAACTCACATTAATTGCGTTGCGCTCACTGCCC
GCTTTCCAGTCGGGAAACCTGTCGTGCCAGCTGCATTAATGAATCGGCCAACGCGCGGGGAG
AGGCGGTTTGCGTATTGGGCGCTCTTCCGCTTCCTCGCTCACTGACTCGCTGCGCTCGGTCG
TTCGGCTGCGGCGAGCGGTATCAGCTCACTCAAAGGCGGTAATACGGTTATCCACAGAATCA
GGGGATAACGCAGGAAAGAACATGTGAGCAAAAGGCCAGCAAAAGGCCAGGAACCGTAAAAA
GGCCGCGTTGCTGGCGTTTTTCCATAGGCTCCGCCCCCCTGACGAGCATCACAAAAATCGAC
GCTCAAGTCAGAGGTGGCGAAACCCGACAGGACTATAAAGATACCAGGCGTTTCCCCCTGGA
AGCTCCCTCGTGCGCTCTCCTGTTCCGACCCTGCCGCTTACCGGATACCTGTCCGCCTTTCT
CCCTTCGGGAAGCGTGGCGCTTTC
```

### HiFi DNA Assembly® Protocol

#### Optimal Quantities

NEB recommends a total of 0.03–0.2 pmols of DNA fragments when 1 or 2 fragments are being assembled into a vector and 0.2–0.5 pmoles of DNA fragments when 4–6 fragments are being assembled. Efficiency of assembly decreases as the number or length of fragments increases. To calculate the number of pmols of each fragment for optimal assembly, based on fragment length and weight, we recommend the following formula:

pmols = (weight in ng) x 1,000 / (base pairs x 650 daltons)

50 ng of 5000 bp dsDNA is about 0.015 pmols.

50 ng of 500 bp dsDNA is about 0.15 pmols.

The mass of each fragment can be measured using the NanoDrop instrument, absorbance at 260 nm or estimated from agarose gel electrophoresis followed by ethidium bromide staining.

#### Assembly Protocol

1. Set up the following reaction on ice:

| | Recommended Amount of Fragments Used for Assembly | | |
|---|---|---|---|
| | 2-3 Fragment Assembly* | 4-6 Fragment Assembly** | Positive Control† |
| Recommended DNA Ratio | vector:insert = 1:2 | vector:insert = 1:1 | |
| Total Amount of Fragments | 0.03–0.2 pmols*<br>X µl | 0.2–0.5 pmols*<br>X µl | 10 µl |
| Assembly Master Mix (2X) | 10 µl | 10 µl | 10 µl |
| Deionized $H_2O$ | 10-X µl | 10-X µl | 0 |
| Total Volume | 20 µl*** | 20 µl*** | 20 µl |

* Optimized cloning efficiency is 50–100 ng of vectors with 2 fold excess of inserts. Use 5 times more inserts if size is less than 200 bps. Total volume of unpurified PCR fragments in the assembly reaction should not exceed 20%.
** To achieve optimal assembly efficiency, it is recommended to design ≥ 20 bp overlap regions between each fragment with equimolarity (suggested: 0.05 pmol each).
† Control reagents are provided for 5 experiments.

‡ If greater numbers of fragments are assembled, increase the volume of the reaction, and use additional Assembly Master Mix.

2. Incubate samples in a thermocycler at 50°C for 15 minutes when 2 or 3 fragments are being assembled or 60 minutes when 4-6 fragments are being assembled. Following incubation, store samples on ice or at –20°C for subsequent transformation.

   Note: Reaction times less than 15 minutes are generally not recommended. Extended incubation times (up to 4 hours) have been shown to improve assembly efficiencies in some cases. Do not incubate the assembly reaction overnight.

3. Transform NEB 5-alpha Competent E. coli cells (provided in the cloning kit or purchased separately from NEB) with 2 µl of the assembled product, following the appropriate transformation protocol.

### Transformation Protocols

**Transformation with chemically competent cells.**

1. Thaw chemically competent cells on ice.

2. Transfer 50 µl of competent cells to a 1.5 ml microcentrifuge tube (if necessary).

3. If the chemically competent cells are from New England Biolabs, add 2 µl of assembled product to NEB competent cells and go to step 4 directly. If competent cells are purchased from other manufacture, dilute assembled products 4-fold with $H_2O$ prior transformation. This can be achieved by mixing 5 µl of assembled products with 15 µl of $H_2O$. Add 2 µl of the diluted assembled product to competent cells.

4. Mix gently by pipetting up and down or flicking the tube 4–5 times. Do not vortex. Place the mixture on ice for 30 minutes. Do not mix.

5. Heat shock at 42°C for 30 seconds.* Do not mix.

6. Transfer tubes on ice for 2 minutes.

7. Add 950 µl of room temperature SOC media* to tubes.

8. Place the tube at 37°C for 60 minutes. Shake vigorously (250 rpm) or rotate.

9. Warm selection plates to 37°C.

10. Spread 100 µl of the cells onto the plates with appropriate antibiotics. Use Amp plates for positive control sample.

11. Incubate plates overnight at 37°C.

    * Please note: Follow the manufacturer's protocols for the duration and temperature of the heat shock step, as well as the optimal medium for recovery. Typically, transformation of our positive control assembly product will yield more than 100 colonies on an Amp plate with greater than 80% colonies containing inserts.

    NEB recommends NEB 5-alpha Competent E. coli (NEB #C2987) for transformation of Gibson Assembly products. It is also possible to use other NEB competent E. coli strains, with the exception of BL21, BL21(DE3), Lemo21(DE3) and Nico21(DE3). For example, Shuffle T7 Express Competent E. coli can be used for the expression of a difficult to express protein. When using competent E. coli from a vendor other than NEB, we have seen decreased robustness of transformation with the Gibson Assembly reaction.

**Transformation with electrocompetent cells.**

1. Thaw electrocompetent cells on ice.
2. Transfer 50 µl of electrocompetent cells to a pre-chilled electroporation cuvette with 1 mM gap.
3. Dilute assembled products 3-fold with $H_2O$ prior electroporation. This can be achieved by mixing 5 µl of assembled products with 10 µl of $H_2O$. Add 1 µl of the diluted assembly product to electrocompetent cells.
4. Mix gently by pipetting up.
5. Once DNA is added to the cells, electroporation can be carried out immediately. It is not necessary to incubate DNA with cells.
6. Add 950 µl of room temperature SOC media to the cuvette immediately after electroporation.
7. Place the tube at 37°C for 60 minutes. Shake vigorously (250 rpm) or rotate.
8. Warm selection plates to 37°C.
9. Spread 100 µl of the cells onto the plates.
10. Incubate overnight at 37°C.
